# From *cereus* to anthrax and back again: The role of the PlcR regulator in the “cross-over” strain *Bacillus cereus* G9241

**DOI:** 10.1101/2022.07.06.498675

**Authors:** Shathviga Manoharan, Grace Taylor-Joyce, Thomas A. Brooker, Carmen Sara Hernandez-Rodriguez, Alexia Hapeshi, Victoria Baldwin, Les Baillie, Petra C. F. Oyston, Nicholas R. Waterfield

## Abstract

*Bacillus cereus* G9241 was isolated from a Louisiana welder suffering from an anthrax-like infection. The organism carries two transcriptional regulators that have previously been proposed to be incompatible with each other: the pleiotropic transcriptional regulator PlcR found in most members of the *Bacillus cereus* group but truncated in all *Bacillus anthracis* isolates, and the anthrax toxin regulator AtxA found in all *B. anthracis* strains and a few *B. cereus* sensu stricto strains. Here we report cytotoxic and haemolytic activity of cell free *B. cereus* G9241 culture supernatants cultured at 25 °C to various eukaryotic cells. However, this is not observed at the mammalian infection relevant temperature 37 °C, behaving much like the supernatants generated by *B. anthracis*. Using a combination of genetic and proteomic approaches to understand this unique phenotype, we identified several PlcR-regulated toxins to be secreted highly at 25 °C compared to 37 °C. Furthermore, we demonstrate that differential expression of the protease involved in processing the PlcR quorum sensing activator molecule PapR appears to be the limiting step for the production of PlcR-regulated toxins at 37 °C, giving rise to the temperature-dependent haemolytic and cytotoxic activity of the culture supernatants. This study provides an insight on how *B. cereus* G9241 is able to ‘switch’ between *B. cereus* and *B. anthracis*–like phenotypes in a temperature-dependent manner, potentially accommodating the activities of both PlcR and AtxA.

## INTRODUCTION

The *Bacillus cereus* sensu lato complex is a group of genetically similar but phenotypically diverse Gram-positive, soil-borne, rod-shaped bacteria (1,2), which includes the well-studied *Bacillus anthracis and Bacillus cereus*. *B. anthracis* is the etiological agent of anthrax (3) while *B. cereus* can colonise hosts as diverse as insects (4) and humans, in which many strains can cause serious foodborne illness (5). Most members of the *B. cereus* group express the chromosomally encoded transcriptional regulator PlcR (<u>P</u>hospho<u>l</u>ipase <u>C</u> <u>r</u>egulator), which controls the expression of many secreted degradative enzymes and toxins (6). However, the *plcR* gene in all *B. anthracis* isolates contains a point mutation, which frameshifts the gene and thus renders it non-functional (7). It has been proposed that the acquisition of AtxA, the mammalian responsive transcriptional regulator involved in expressing anthrax toxins, is incompatible with the activity of PlcR, leading to a selection for PlcR mutation and inactivation (7,8). Interestingly, a *B. cereus-B. anthracis* “cross-over” strain designated *B. cereus* G9241 (hereon referred to as *Bc*G9241) encodes intact copies of both *atxA* and *plcR* genes (9), suggesting this incompatibility dogma is not as straightforward as first suggested.

*Bc*G9241 was isolated from a Louisiana welder, who was hospitalised with a respiratory infection resulting in a case of potentially lethal pneumonia (9). Symptoms were similar to those of inhalational anthrax. The patient also suffered with haemoptysis. *Bc*G9241 possesses three extrachromosomal elements: pBCX01, pBC210 and pBFH_1 (9,10). The plasmid pBCX01 shares 99.6% sequence homology with the plasmid pXO1 from *B. anthracis* strains. pBCX01 encodes the protective antigen (PA), lethal factor (LF), oedema factor (EF) and the AtxA1 regulator. The second plasmid pBC210 (previously known as pBC218) encodes for the *B. cereus* exo-polysaccharide (BPS) capsule biosynthesis genes, *bpsXABCDEFGH* (9). A novel toxin named certhrax is also encoded on the pBC210 plasmid (11,12), which has 31% amino acid sequence similarity with the LF from *B. anthracis*. Moreover, pBC210 encodes gene products with amino acid sequences bearing homology to AtxA and PA of *B. anthracis* (9). Subsequently these genes have been named *atxA2* and *pagA2*. The third extrachromosomal element pBFH_1 (previously known as pBClin29) is a linear phagemid (9). Although the sequence is available for pBFH_1, it is not known if it contributes to the lifestyle of *Bc*G9241. Our group demonstrated by transmission electron microscopy that the pBFH_1 phage could be produced and released into the supernatant (13). The shape of the phage particles and the dimensions of the tail and head appeared to be consistent with the Siphoviridae family (14), suggesting the pBFH_1 is a Siphoviridae phage. Phenotypically, *Bc*G9241 is haemolytic and resistant to γ-phage like other *B. cereus* strains (9). Further phenotypic and genetic analyses suggested that *Bc*G9241 should be considered a member of the *B. cereus* sensu stricto group as it does not encode a point mutation in the *plcR* gene indicative of a *B. anthracis* strain (8).

PlcR controls the expression of many secreted enzymes and toxins (6,7,15), with at least 45 regulated genes found in *B. cereus* type strain ATCC 14579 (16), hereon referred to as *Bc*ATCC14579. These secreted proteins, which contribute significantly to virulence in mice and insects (17,18), include haemolysins, enterotoxins, proteases, collagenases and phospholipases (15). Activation of PlcR requires the binding of a secreted, processed and reimported form of the signalling peptide PapR (6,19–21). The *papR* gene is located downstream of *plcR* and encodes a 48-amino acid protein. PapR48 is secreted from the cell via the Sec machinery and processed to a heptapeptide by the extracellular zinc metalloprotease, NprB and potentially other extracellular proteases (22). The *nprB* gene is often tightly linked to the *plcR-papR* operon, but in the opposite orientation (6,21,22). The processed form PapR_7_ is reimported into the bacterium by the oligopeptide permease (Opp) system (23). The processed form of PapR can then bind and activate PlcR. The active PlcR-PapR complex binds to the palindromic operator sequence (PlcR box: TATGNAN4TNCATA) found in the promoter regions of the regulon genes, subsequently activating transcription of these genes (24–26). PlcR also positively auto-regulates its own transcription, which can be repressed by the sporulation factor Spo0A, facilitated by two Spo0A boxes flanking the PlcR box (27). Four distinct classes of PlcR-PapR systems have evolved and differ by the 5 C-terminal amino acids of PapR, which bind to PlcR, with PapR from one group unable to activate the transcriptional activity of PlcR from another (28).

The chromosome of *Bc*G9241 encodes a large range of intact exotoxin genes confirming the strain is part of the sensu stricto group (9). Several of the toxin genes are likely to be regulated by PlcR, by virtue of the presence of the PlcR-box sequence in the promoter regions (6). These include haemolysin BL (Hbl) encoded by *hblCDAB*, the tripartite non-haemolytic enterotoxin (Nhe), encoded by *nheABC*, and the enterotoxin cytotoxin K (CytK), encoded by *cytK*. These three toxins are all classed as enterotoxins and have been isolated from patients suffering from food-borne, diarrhoeal infections (29–31). Since isolating *Bc*G9241, cases of anthrax-like disease caused by other non-*B. anthracis* bacteria have been reported, affecting both animals such as chimpanzees and gorillas in the Ivory Coast and Cameroon during the early 2000s (32–36), in addition to humans (9,10,37–44). Some of these isolates carry functional copies of *plcR* and *atxA* (recently reviewed in (45)); this warrants further investigation into the role of PlcR in these strains, as the loss of PlcR activity has been proposed to be crucial in anthrax disease caused by *B. anthracis*. So far, only one study on *Bc*G9241 has been carried out to identify how PlcR, AtxA and their respective regulons are expressed. A microarray assay carried out by (46) demonstrated that in *Bc*G9241 the *plcR* gene was ~2.4 fold more highly expressed in an aerobic environment compared to when exposed to CO_2_/bicarbonate, while in contrast, the *atxA1* gene showed higher expression in CO_2_ by ~5.6 fold. Understanding how the PlcR-PapR regulatory circuit acts in *B. cereus-B. anthracis* “cross-over” strains may provide an insight into their evolution and give a more complete picture of the phylogeny.

Here, we describe the temperature-dependent haemolytic and cytolytic activity of *Bc*G9241, caused by PlcR-controlled toxins and proteases. We also identify the limiting step in the PlcR-PapR circuit involved in preventing the expression of PlcR-regulated toxins at 37 °C. NprB is not involved in processing PapR in *Bc*G9241 and other *B. cereus* strains carrying functional copies of both *plcR* and *atxA*. We hypothesise that a change in the PlcR-PapR regulatory network in *Bc*G9241 may have allowed the carriage of intact copies of both *plcR* and *atxA*, by virtue of a temperature-dependent suppression of the PlcR-PapR circuit and the loss of the *nprB* gene.

## RESULTS

### *Bc*G9241 culture supernatants demonstrate temperature-dependent toxicity against a range of eukaryotic cells

It has been previously shown in other members of the *B. cereus* group that PlcR regulates the secretion of multiple virulence proteins, such as cytolytic toxins and enzymes involved in macromolecule degradation (16,47,48). We therefore tested the haemolytic activity of cell free culture supernatants from *Bc*G9241 cultures grown at 25 °C and 37 °C to sheep erythrocytes. We tested the effect of growth at 25 °C and 37 °C to partially emulate environmental and mammalian host conditions, respectively. Filtered supernatants were extracted from cultures grown to exponential phase (OD_600_=0.5) and stationary phase (OD_600_=1.5). From 25 °C cultures, the supernatants from exponential and stationary phases demonstrated haemolytic activity to red blood cells (RBCs), above the 75% level by comparison to the expected lysis from the positive control (1% Triton X100) (**Fig 1**). In contrast, supernatants from *Bc*G9241 grown at 37 °C showed very little lytic activity (**Fig 1**). This led us to the hypothesis that *Bc*G9241 ‘switches’ its phenotype from a haemolytic *B. cereus*-like phenotype at 25 °C to a non-haemolytic *B. anthracis*-like phenotype at 37 °C.

**Figure 1:**
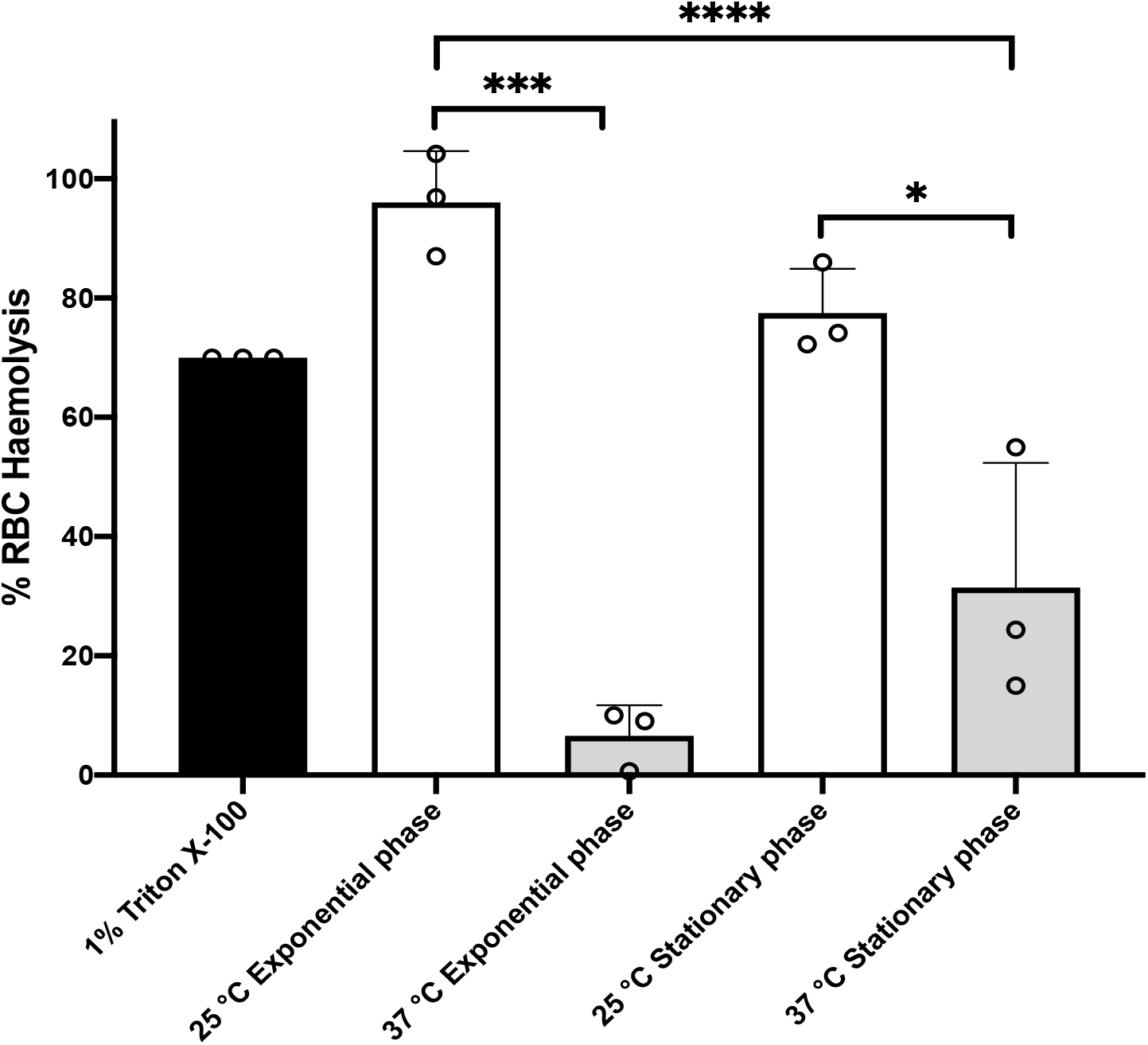
*Bc*G9241 supernatant is significantly more toxic to sheep RBCs, when extracted from a 25 °C grown culture compared to a 37 °C grown culture. The haemolysis assay was conducted by incubating *Bc*G9241 supernatant with 4% RBCs for 1 hour at 37 °C. The OD_540_ was measured, and RBC lysis was calculated as a % of expected RBC lysis by Triton X-100 (1% v/v). Stars above columns represent significance levels. * denotes an unpaired t test with a p-value of 0.0232; *** denotes a Welch’s t test with a p-value of 0.0003; **** denotes an ordinary one-way ANOVA with a p-value of <0.0001. Error bars denote one standard deviation, and all samples were to an n=3.

We expanded the study to test the toxicity of cell free culture supernatants from a range of *Bacillus* cultures grown at 25 °C against *ex vivo Manduca sexta* haemocytes. Filtered supernatants were extracted from cultures grown for 16 h. Microscopic examination showed that supernatants from the reference strain *Bc*ATCC14579, *B*cG9241 and *Bc*G9241 ΔpBCX01 (in which the plasmid had been cured) caused extensive lysis of the *M. sexta* haemocytes (**Fig S1A**). In contrast, supernatants from *Bacillus thuringiensis* 407 Cry^−^ *ΔplcR* (*Bt ΔplcR*) mutant strain were innocuous, showing no difference from the negative buffer control (**Fig S1A**). *Bt ΔplcR* is an accepted Δ*plcR B. cereus* model as the crystal toxin plasmid has been cured (27). The cytotoxicity observed with *Bc*G9241 and *Bc*G9241 ΔpBCX01 indicated that cytotoxins were secreted by both strains and is possibly unaffected by the presence of pBCX01 plasmid at 25 °C (**Fig S1A**).

We quantified the effect of supernatants from these same strains using haemocyte cell viability assays. We also expanded the study to include supernatants from the *B. anthracis* Sterne strain, which lacks the pXO2 plasmid (hereon referred to as *Ba* St). Like all *B. anthracis* strains, *Ba* St has a frame-shifted copy of the *plcR* gene. From 25 °C grown cultures, we observed cytotoxicity responses consistent with the microscopic examinations (**Fig S1B**), with the supernatants of *Bc*ATCC14579, *Bc*G9241 and *Bc*G9241 ΔpBCX01, all showing potent toxicity. In contrast, supernatants from *Ba* St and *Bt ΔplcR* showed little or no cytotoxicity (**Fig S1B**). However, when grown at 37 °C, cytotoxicity of *Bc*G9241 and BcG9241 ΔpBCX01 supernatants was highly attenuated, to levels no different from those of the *Ba* St and *Bt ΔplcR* supernatants (**Fig S1C**).Cytotoxicity of the *Bc*ATCC14579 supernatant was still observed at 37 °C (**Fig S1C**). Temperature-dependent cytotoxic activity of *Bc*G9241 and *Bc*G9241 ΔpBCX01 supernatants were also observed in a range of mammalian cells including T2-lymphocytes, polymorphonuclear leukocytes and macrophages (using supernatant extracted from *Bacillus* cultures grown for 16 h), which re-capitulated the trend seen with the insect haemocytes (**Fig S2**).

### Temperature and growth phase-dependent proteomic analysis of *Bc*G9241 culture supernatants

In order to investigate the potential cytolytic and haemolytic factors secreted by *Bc*G9241, we analysed the proteomic profiles of supernatants from cultures grown at 25 °C and 37 °C in LB broth, taken from both mid-exponential (OD_600_ = 0.5) and stationary growth phases. For stationary phase, *Bc*G9241 cultures supernatant were extracted after 10 hours growth at 25 °C and after 7 hours growth at 37°C (13). Proteins were run through nanoLC-ESI-MS and peptide reads were counted using MaxQuant (Max Planck Institute). Comparisons were made using the Perseus software (Max Planck Institute) and plotted as the difference in proteins expressed between the two temperatures. The full datasets generated can be seen in the **Supplementary Dataset S1 and S2**.

A principal component analysis (PCA) was generated to show the variance between all biological replicates of the *Bc*G9241 supernatants collected from both exponential- and stationary phases. The PCA plots revealed that protein extracts from the exponential phase supernatants overlap with each other, not forming distinct clusters and are highly reproducible (**Fig S3**). The plot also showed that growth temperature affected the protein profiles more significantly at stationary phase compared to exponential phase (**Fig S3**). Furthermore, protein profiles extracted from stationary phase growth at 37 °C were more variable than those from other conditions (**Fig S3**).

#### A diverse and abundant toxin “profile” was secreted at 25 °C, while high levels of phage proteins were secreted at 37 °C during exponential growth phase of *Bc*G9241

With the cut-off criteria of p-value < 0.05 and a minimum 2-fold change in protein level, 33 supernatant proteins were identified as being significantly more abundant at 25 °C compared to 37 °C. Of these, 11 of the 12 most highly expressed are known toxin homologs (**Table 1** and **Fig S4**). This included all components of the Hbl toxin encoded by the *hbl* operon AQ16_4930 – 4933 **(Fig S4-purple arrows)**. Other known cytotoxic proteins were also abundant in the supernatant at 25 °C compared to 37 °C, including the Nhe toxin encoded by the *nhe* operon AQ16_658 – 660 (**Fig S4-green arrows**), a collagenase (AQ16_1941), a thermolysin metallopeptidase (AQ16_5317), phospholipase C (Plc, AQ16_1823) and CytK (AQ16_1392).

**Table 1:** The 15 most abundant toxins at higher levels at the two temperature in the secretome of *Bc*G9241 during exponential growth and stationary phase. The significance cut-off criteria used was a p-value of <0.05 and a minimum of a 2-fold change in protein level.

| Log<sub>2</sub>-Fold Change | 25 °C > 37 °C at exponential phase | Gene | Gene Loci (AQ16_) |
| --- | --- | --- | --- |
| 6.46 | Haemolysin BL lytic component L2 |  | 4931 |
| 5.88 | Non-haemolytic enterotoxin binding component | *nheC* | 658 |
| 5.67 | Hemolysin BL-binding component | *hblA* | 4932 |
| 4.41 | Collagenase family protein |  | 1941 |
| 4.31 | Extracellular ribonuclease | *bsn* | 4754 |
| 3.98 | Hemolysin BL-binding component | *hblB* | 4933 |
| 3.93 | Non-hemolytic enterotoxin lytic component L2 | *nheA* | 660 |
| 3.84 | Thermolysin metalloproteinase, catalytic domain protein |  | 5317 |
| 3.77 | Non-hemolytic enterotoxin lytic component L1 | *nheB* | 659 |
| 3.57 | Haemolysin BL lytic component L1 |  | 4930 |
| 3.20 | Phospholipase C | *olc* | 1823 |
| 2.69 | Cytotoxin K | *cytK* | 1392 |
| 2.49 | THUMP domain protein |  | 931 |
| 2.44 | Probable butyrate kinase | *buk* | 3880 |
| 2.42 | DEAD-box ATP-dependent RNA helicase | *cshA* | 2258 |
| **37 °C > 25 °C at exponential phase** |  |  |  |
| 5.27 | Phage family protein | *gp49* | 5822 |
| 4.96 | Putative phage major capsid protein | *gp34* | 5824 |
| 4.53 | Prophage minor structural protein |  | 5836 |
| 4.32 | Putative gp14-like protein | *gp14* | 5832 |
| 4.31 | N-acetylmuramoyl-L-alanine amidase family protein |  | 5839 |
| 3.65 | Phage tail family protein |  | 5835 |
| 3.40 | Putative major capsid protein | *gpP* | 5831 |
| 2.89 | Uncharacterized protein |  | 5823 |
| 2.36 | WxL domain surface cell wall-binding family protein |  | 3215 |
| 2.34 | WxL domain surface cell wall-binding family protein |  | 3217 |
| 2.33 | Phage antirepressor KilAC domain protein |  | 5855 |
| 2.30 | Dihydropteroate synthase | *folP* | 2448 |
| 2.19 | Zinc-binding dehydrogenase family protein |  | 318 |
| 2.07 | WxL domain surface cell wall-binding family protein |  | 3218 |
| 1.95 | Toxic anion resistance family protein |  | 2068 |
| **25 °C > 37 °C at stationary phase** |  |  |  |
| 7.83 | Thermolysin metalloproteinase, catalytic domain protein |  | 5317 |
| 4.86 | Transglutaminase-like superfamily protein |  | 1487 |
| 4.54 | UDP-N-acetylglucosamine 1-carboxyvinyltransferase | *murA* | 2685 |
| 4.51 | Ribonuclease J | *rnjA* | 2375 |
| 4.51 | Ornithine aminotransferase | *rocD* | 1349 |
| 4.16 | Pyruvate carboxylase | *pyc* | 4104 |
| 4.10 | Malic enzyme, NAD binding domain protein |  | 3400 |
| 4.01 | CTP synthase | *pyrG* | 2681 |
| 3.95 | Glycerophosphoryl diester phosphodiesterase family protein |  | 4572 |
| 3.73 | Leucine--tRNA ligase | *leuS* | 3254 |
| 3.66 | Viral enhancin family protein |  | 2918 |
| 3.66 | Subtilase family protein |  | 4301 |
| 3.16 | 50S ribosomal protein L2 | *rplB* | 2391 |
| 3.16 | Aldo/keto reductase family protein |  | 2308 |
| 3.14 | Sphingomyelin phosphodiesterase | *sph* | 1822 |
| **37 °C > 25 °C at stationary phase** |  |  |  |
| 9.19 | Chitinase A1 | *chiA1* | 2089 |
| 7.85 | Putative hydrolase |  | 2662 |
| 7.31 | Glucanase |  | 5335 |
| 6.99 | Collagenase family protein |  | 4546 |
| 5.67 | Chitinase A |  | 4342 |
| 4.83 | Peptide ABC transporter |  | 2309 |
| 4.54 | Calcineurin-like phosphoesterase family protein |  | 4913 |
| 4.43 | Urocanate hydratase | *hutU* | 4415 |
| 4.31 | Single-stranded DNA-binding protein | *ssb* | 2546 |
| 4.28 | Matrixin family protein |  | 4915 |
| 3.77 | 3-hydroxyacyl-[acyl-carrier-protein] dehydratase FabZ | *fabZ* | 2750 |
| 3.56 | Formate acetyltransferase | *pflB* | 2025 |
| 3.52 | Ribose-phosphate pyrophosphokinase | *prs* | 2472 |
| 3.26 | Putative phage major capsid protein |  | 5824 |
| 3.22 | 50S ribosomal protein L4 | *rplD* | 2393 |

Conversely, at 37 °C, the secretome contained negligible levels of these cytotoxic proteins (if present at all). There was, however, an abundance of phage capsid proteins encoded by the pBFH_1 phagemid at 37 °C compared to 25 °C (**Fig S4-black arrows**). More specifically 25 proteins were found to be more abundant in the secretome at 37 °C compared to 25 °C. The 10 most abundant proteins at 37 °C compared to 25 °C were encoded by the pBFH_1 phagemid (**Table 1**). Proteins from an operon of WxL-domain cell wall-binding proteins were also seen to be more abundant at 37 °C compared to 25 °C (AQ16_3217 – 3219).

#### The temperature dependent *Bc*G9241 secretome at stationary growth phase

Between 25 °C and 37 °C, 51 proteins showed temperature dependent differences (**Fig 2B**). Unlike the mid-exponential observations, the more abundant proteins in the 25 °C stationary phase supernatants were not all cytotoxins, although several enzymes were present (**Table 1**). In fact, of the 11 toxins seen to be more abundant at 25 °C during exponential phase growth, only AQ16_5317 was identified at higher levels at 25 °C during stationary phase. This is a thermolysin metallopeptidase, which has a PlcR-box present in the promoter region (**Table S1**), and is over 200-fold more abundant at 25 °C. The relevance of this is discussed below. Several of the more abundant proteins (e.g. AQ16_3254, 4226, 374) identified were likely cellular proteins, possibly indicating greater autolysis at 25 °C compared to 37 °C. The top 5 proteins more abundant in 37 °C compared to 25 °C supernatants were all extracellular enzymes including two chitinases, a hydrolase, a glucanase and a collagenase (**Table 1**). In addition, a matrixin family protein (AQ16_4915), another extracellular enzyme, was also identified as 4.3 log2-fold higher at 37 °C. Again, we saw cellular components including 50S ribosome subunit proteins and RecA, which possibly signified cell lysis. Only one of the phage capsid proteins identified as higher at 37 °C in the exponential phase secretome, Gp34 (AQ16_5824), was significantly higher at 37 °C in stationary phase.

**Figure 2:**
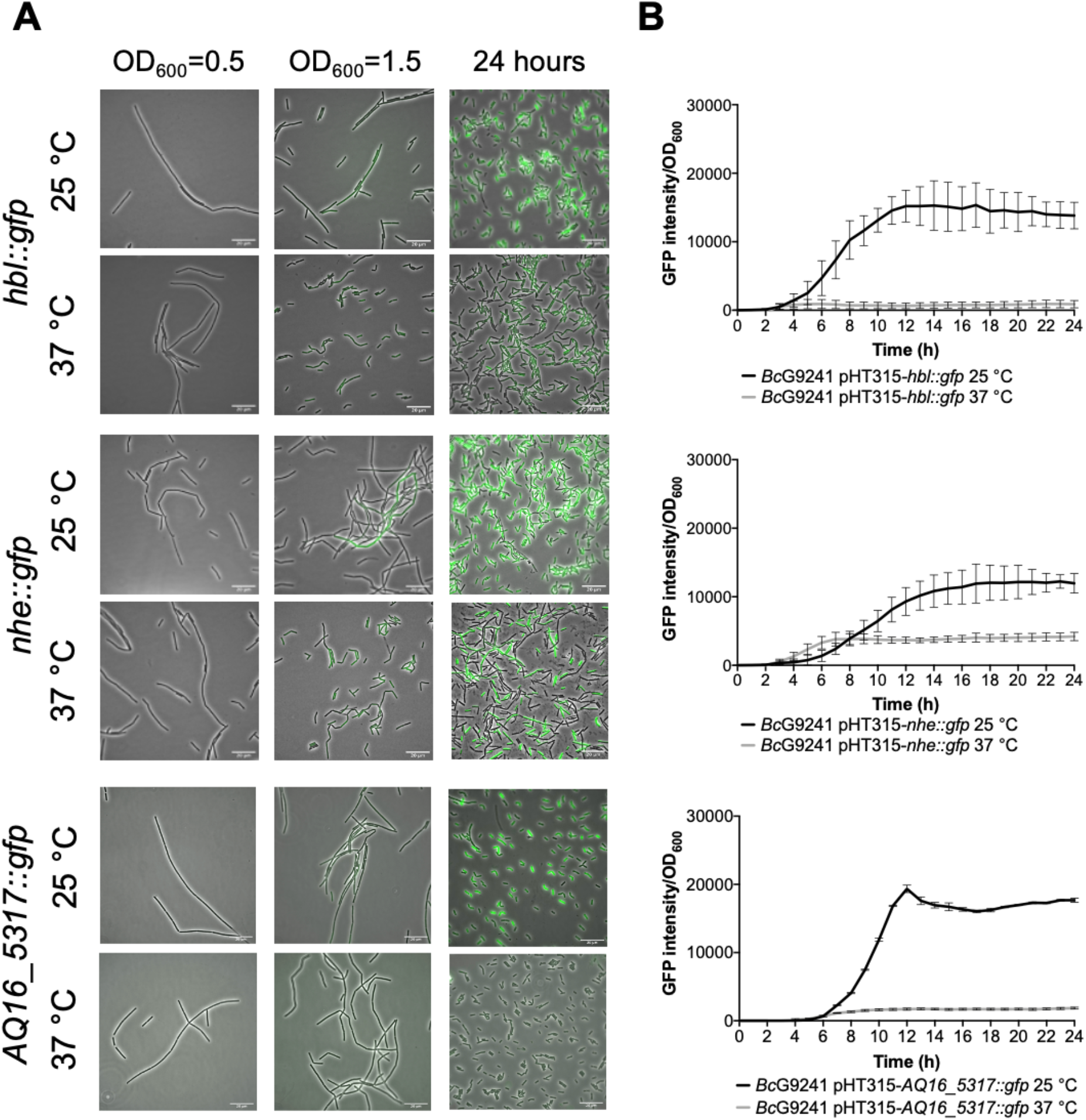
Temperature dependent expression of PlcR-regulated toxins and enzymes in *Bc*G9241 using GFP reporters. (**A**) A representative selection of microscopy images of the transcription-translation GFP reporters of PlcR-regulated toxins for *Bc*G9241 taken at three different time points: mid-exponential phase which is 2 hours at 37 °C and 5 hours at 25 °C (OD_600_=0.5), early stationary phase which is 4 hours at 37 °C and 7 hours at 25 °C (OD_600_=1.5) and 24 hours. (**B**) **Fluorescence of toxin reporters over time in LB**. GFP intensity/OD_600_ and change in GFP (ΔGFP/OD_600_) of *Bc*G9241 containing PlcR-regulated toxin reporters over 24 hours growth in 100 μl volume LB media at 25 °C (in black) and 37 °C (in grey). Each line represents the mean of three biological replicates with three technical replicates each and error bars denote standard deviation.

### Temperature-dependent cell proteome analysis of exponentially growing *Bc*G9241 cells

The greatest temperature-dependent change in secreted toxin profiles was seen in exponentially growing cells. Therefore, to investigate the potential role of PlcR in the temperature-dependent regulation of toxin secretion, and any relationship between protein synthesis and secretion, a proteomic analysis of whole cells was performed. The same samples used for the supernatant proteomic analysis were used for this, allowing for direct correlation of the datasets. The full datasets generated can be seen in the **Supplementary Dataset S3.**

#### No build-up of toxins was observed in the cellular proteome of *Bc*G9241 at 37 °C exponential phase

With a cut-off criteria of p-value < 0.05 and a minimum 2-fold change in protein level, 67 proteins were found to be significantly more abundant at 25 °C compared to 37 °C. The most abundant proteins at 25 °C compared to 37 °C included cold shock proteins CspA and YdoJ family proteins (**Table 2** and **Fig S5**). Only two of the toxin proteins seen at higher levels at 25 °C in comparison to 37 °C in the secretome were also significantly higher in the cell proteome, NheA and NheB (AQ16_659 and 660).

**Table 2:** Top 15 cellular proteins that are more abundant at each temperature in exponentially growing *Bc*G9241. The significance cut-off criteria used was a p-value of <0.05 and a minimum 2-fold change in protein level.

| Log<sub>2</sub>-Fold Change | 25 °C > 37 °C at exponential phase | Gene | Gene Loci (AQ16_) |
| --- | --- | --- | --- |
| 5.09 | Major cold shock protein | *cspA* | 1368 |
| 4.72 | Uncharacterized protein |  | 4251 |
| 4.48 | Cold-inducible YdjO family protein |  | 175 |
| 4.10 | Uncharacterized protein |  | 4821 |
| 3.18 | Transglutaminase-like superfamily protein |  | 1487 |
| 3.06 | Flagellar motor switch FlIM family protein |  | 858 |
| 2.77 | FMN-dependent NADH-azoreductase | *azoR4* | 2611 |
| 2.57 | Uncharacterized protein |  | 1372 |
| 2.47 | Hemolytic enterotoxin family protein |  | 659 |
| 2.41 | Uncharacterized protein |  | 1559 |
| 2.21 | Major cold shock protein | *cspA* | 174 |
| 2.20 | SET domain protein |  | 2908 |
| 2.19 | Transposase family protein |  | 1725 / 4355 |
| 2.15 | Rhodanese-like domain protein |  | 1704 |
| 2.09 | Hemolytic enterotoxin family protein | *nheA* | 660 |
| **37 °C > 25 °C at exponential phase** |  |  |  |
| 4.34 | WxL domain surface cell wall-binding family protein |  | 3218 |
| 3.74 | Uncharacterized protein |  | 3219 |
| 2.87 | Formate acetyltransferase | *pflB* | 2025 |
| 2.73 | Uncharacterized protein |  | 1429 |
| 2.29 | DNA repair protein | *recN* | 3857 |
| 2.27 | DNA protection during starvation protein 1 | *dps1* | 512 |
| 2.25 | Uncharacterized protein |  | 5765 |
| 2.25 | Pyruvate formate-lyase-activating enzyme | *pflA* | 2024 |
| 2.23 | L-lactate dehydrogenase | *ldh* | 3111 |
| 2.166 | CamS sex pheromone cAM373 family protein |  | 2171 |
| 2.09 | L-asparaginase, type I family protein |  | 4939 |
| 1.98 | Heat-inducible transcription repressor | *hrcA* | 3712 |
| 1.97 | Uncharacterized protein |  | 5849 |
| 1.96 | Membrane MotB of proton-channel complex MotA/MotB family protein |  | 3490 |
| 1.84 | Periplasmic binding family protein |  | 1888 |

51 proteins were found to be significantly more abundant at 37 °C compared to 25 °C (**Table 2**). Proteins from an operon of WxL-domain cell wall-binding proteins were seen to be more abundant at 37 °C (AQ16_3217 – 3219). In addition, various heat stress response proteins were also identified as higher at 37 °C. These include: AQ16_3857, a DNA repair protein; AQ16_512, a DNA protection protein and a thermosensor operon, AQ16_3712 – 3714, involved in protein refolding. Interestingly, despite the significantly increased abundance in the secretome, only two proteins encoded on the pBFH_1 phagemid (AQ16_5849 and _5858) showed increased abundance in the cell proteome, both of which are uncharacterised.

A build-up of toxins from the cell proteome at 37 °C was not observed, demonstrating that temperature-dependent toxin expression is not regulated at the level of secretion. PlcR was detected at both temperatures with no significant difference in abundance levels.

### Analysis of PlcR-controlled toxin expression in *Bc*G9241

Haemolytic and cytolytic assays have suggested that *Bc*G9241 containing a functional copy of the *plcR* gene show temperature-dependent toxicity. Hbl, Nhe, Plc, CytK and a thermolysin metallopeptidase (AQ16_5317), which are regulated by the PlcR-PapR circuit (16), were detected with high abundance in the secretome analysis at 25 °C compared to 37 °C. In order to confirm the temperature-dependent toxin and protease production, a panel of transcription-translation reporter plasmids were made, in which the promoter regions and the first 24 bp of the coding sequence of *hbl, nhe, plc, cytK* and *AQ16_5317* were genetically fused in frame to a *gfp* gene with no start codon (referred to hereon as *hbl∷gfp, nhe∷gfp, plc∷gfp, cytK∷gfp* and *AQ16_5317∷gfp*). Note that only eight N-terminal amino acids from the ORF were cloned as it is not sufficient to serve as a Sec-dependant secretion signal for the toxins, preventing the GFP from being secreted. For comparison, GFP reporters of PlcR-regulated toxins were also constructed for *Bc*ATCC14579 from homologous regions. Each of the reporter constructs were then transformed into the relevant *B. cereus* strain and examined using fluorescence microscopy and microtitre plate reader assays to assess the expression patterns across growth phases at 25 °C and 3 °C, when grown in LB while maintaining plasmid marker selection. The rate of change in fluorescence (ΔGFP/OD_600_) was calculated every hour by subtracting the fluorescence at a given time point by the fluorescence of the previous time point. This would reveal when the biggest change in GFP expression occurs across the growth phase.

From the microscopy images, the expression of the toxin reporters in *Bc*G9241 was not detected during mid-exponential phase at 25 °C and 37 °C (**Fig 2A** and **Fig S6**). However, by quantifying the GFP intensity of *B. cereus* strains containing the reporters, there were cells with higher fluorescence compared to the control cells (being above the threshold), suggesting that GFP, and therefore the PlcR-regulated proteins were being expressed. The mean GFP intensity of individual cells quantified was higher at 25 °C compared to 37 °C for *hbl∷gfp*, *nhe∷gfp* and *cytK∷gfp* (**Fig S8**). Once reaching stationary phase, the difference in the expression of the toxin reporters in *Bc*G9241 was more pronounced between 25 °C and 37 °C (**Fig 2A** and **Fig S6**). From GFP intensity quantification of individual cells from the micrographs, the mean GFP intensity of *hbl∷gfp, nhe∷gfp, plc∷gfp* and *AQ16_5317∷gfp* at the onset of stationary phase and 24 hours was higher at 25 °C compared to 37 °C in *Bc*G9241 (**Fig S7**). It is interesting to note that *Bc*G9241 cells formed filamentous-like structures during exponential phase which reverted to shorter vegetative rod morphologies once stationary phase was reached.

When the GFP intensity/OD_600_ of *Bc*G9241 harbouring the PlcR-regulated toxin reporters was monitored with a microtitre plate reader over 24 hours (**Fig 2B** and **Fig S6**), the GFP expression was greater at 25 °C compared to 37 °C for *hbl∷gfp, nhe∷gfp, plc∷gfp* and *AQ16_5317∷gfp* while the expression of *cytK∷gfp* appeared similar between both temperatures. By calculating the rate of change in fluorescence (ΔGFP), a large and broad ΔGFP peak was observed at 25 °C while the peak appeared tighter at 37 °C for *hbl∷gfp, nhe∷gfp, plc∷gfp* and *AQ16_5317∷gfp* (**Fig S8**). In comparison, expression of *hbl∷gfp*, *nhe∷gfp* and *BC_2735∷gfp* (BC_2735 has a 97% identity to AQ16_5317 using BLASTP) in *Bc*ATCC14579 appeared to be similar at 25 °C and 37 °C, while the expression of *plc∷gfp* and *cytK∷gfp* was temperature-dependent (**Fig S9**).

### Population level analysis of PlcR and PapR expression in *Bc*G9241

Subsequently, we expanded the analysis by including a panel of transcription-translation reporter plasmids for PlcR and PapR in *Bc*G9241 and *Bc*ATCC14579. The promoter regions and the first 24 bp of the coding sequence of *plcR* and *papR* were genetically fused in frame to a *gfp* gene with no start codon (referred to hereon as *plcR∷gfp* and *papR∷gfp*). Note that only eight N-terminal amino acids from the ORF were cloned as it is not sufficient to serve as a Sec-dependent secretion signal for PapR and to make sure that the PlcR protein was not interfering with the GFP protein. Each of the reporter constructs were then transformed into the relevant *B. cereus* strain and examined using fluorescence microscopy to assess the expression patterns across growth phases at 25 °C and 37 °C, when grown in LB and maintaining plasmid marker selection.

#### Expression of PlcR is not temperature dependent

Expression of *plcR∷gfp* was first observed during early stationary phase at 25 °C and 37 °C (**Fig 3A**). By 24 hours, levels of *plcR∷gfp* increased at both temperatures, with a high level of population heterogeneity in expression within the cell population. The cells that expressed *plcR∷gfp*, also did so at a high level. As observed in *Bc*G9241, expression of *plcR∷gfp* in *Bc*ATCC14579 was also heterogeneous within the cell population (**Fig 3A**). Image analysis provided an objective quantification of this heterogeneous expression observed within the cell population, with a small number of cells expressing *plcR∷gfp* in both *B. cereus* strains (**Fig S10**).

**Figure 3:**
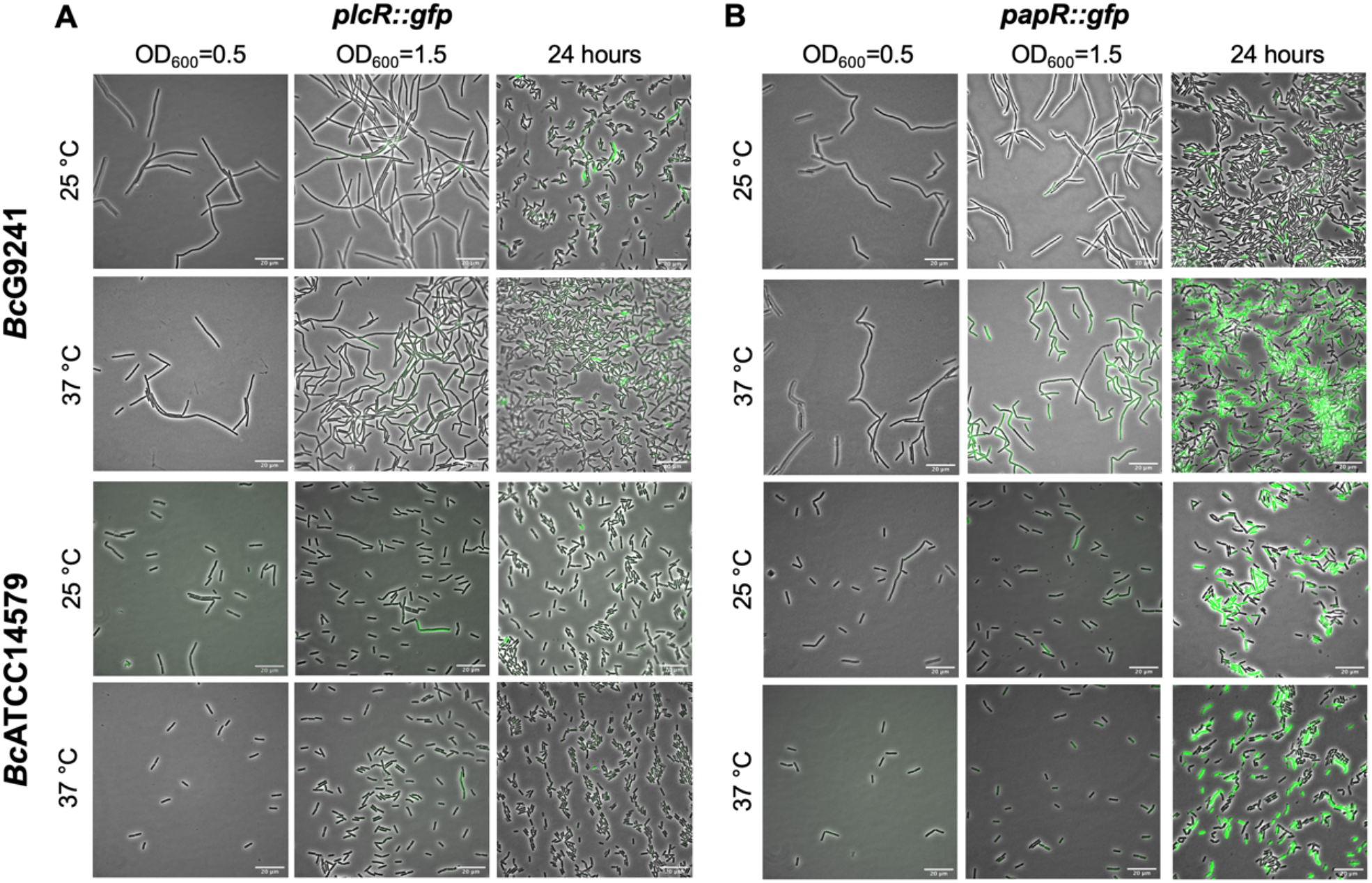
A representative selection of microscopy images of BcG9241 and BcATCC14579 harbouring the transcription-translation GFP reporters of *plcR∷gfp* and *papR∷gfp*. Micrographs were taken at three different time points: mid-exponential phase which is 2 hours at 37 °C and 5 hours at 25 °C (OD_600_=0.5), early stationary phase which is 4 hours at 37 °C and 7 hours at 25 °C (OD_600_=1.5) and 24 hours. Scale bar = 20 *μ*m.

There is a possibility that the population heterogeneity observed for the expression of *plcR∷gfp* could be due to cell death or an error from the reporter itself. To determine whether the *plcR∷gfp* expression was indeed originated from a minority of cells, some potential issues were analysed. The shuttle vector used for the reporter pHT315 encodes an erythromycin resistance gene, and therefore the antibiotic was added to maintain selection. To rule out heterogeneity due to cell death, propidium iodide staining was carried out. Cell viability is assessed when propidium iodide penetrates damaged membranes binding to nucleic acid, leading to fluorescence. At early stationary phase, only a few cells were stained by propidium iodide while within a large population of live cells, a small proportion of cells expressed *plcR∷gfp* (**Fig S11**). This confirms the heterogeneous expression of *plcR∷gfp* was indeed originated from a small subpopulation of live cells and this is not a consequence of cell death in any non-reporter expressing cells.

#### PapR in *Bc*G9241 is highly expressed at 37 °C compared to 25 °C

Expression of *papR∷gfp* was first noticed during early stationary phase at 25 °C and 37 °C. Levels of *papR∷gfp* expression greatly increased by 24 hours, with stronger fluorescence observed by microscopy at 37 °C compared to 25 °C (**Fig 3B**). Image analysis has provided an assessment of the expression observed within the cell population, with a sub-population of cells expressing *papR∷gfp* at 25 °C. In *Bc*G9241 and *Bc*ATCC14579, the mean GFP expression was higher at 37 °C compared to 25 °C (**Fig S10**).

### The import of mature PapR_7_ is functional at 25 °C and 37 °C in *Bc*G9241

A build-up of toxin proteins in the cell proteome at 37 °C was not detected, suggesting that temperature-dependent toxin expression is not regulated at the level of secretion. This led us to investigate whether the import of mature PapR is not functional at 37 °C, causing the temperature-dependent haemolysis and cytolysis phenotypes observed in *Bc*G9241. To understand whether the import system is functional at 37 °C, a haemolysis assay was carried out using supernatants of *B. cereus* cultures grown at 25 °C and 37 °C with mature synthetic PapR peptides added exogenously. Previous studies have demonstrated that the heptapeptide PapR_7_ is the mature form of the quorum sensing peptide (22,28) and therefore synthetic peptides of this form (G9241 PapR_7_= SDLPFEH, ATCC14579 PapR_7_= KDLPFEY) were used.

At 25 °C, with the addition of exogenous self PapR_7_ (i.e., adding G9241 PapR_7_ into cultures of *Bc*G9241 or adding ATCC14579 PapR_7_ into cultures of *Bc*ATCC14579), no significant change in haemolytic activity of the mid-exponential supernatants of *Bc*G9241 was observed, compared to the absence of exogeneous PapR_7_. Nevertheless, haemolytic activity was still observed with/without the addition of the cognate PapR_7_ from supernatants collected at 25 °C in both *Bc*G9241 and *Bc*ATCC14579 (**Fig 4**). In comparison, upon the addition of exogenous PapR a significant change in haemolytic activity of the mid-exponential supernatant of *Bc*ATCC14579 was observed, compared to the absence of exogeneous PapR_7_. At 37 °C, with the addition of cognate PapR_7_, there was a significant increase in haemolytic activity with the mid-exponential and stationary phase *Bc*G9241 supernatant (**Fig 4)**. This suggests that PapR_7_ can get taken into the cell through an import system at 37 °C. In comparison, upon the addition of exogenous cognate PapR_7_, no significant change in haemolytic activity of the mid-exponential supernatant of *Bc*ATCC14579 was observed, compared to the absence of exogeneous PapR_7_.

**Figure 4:**
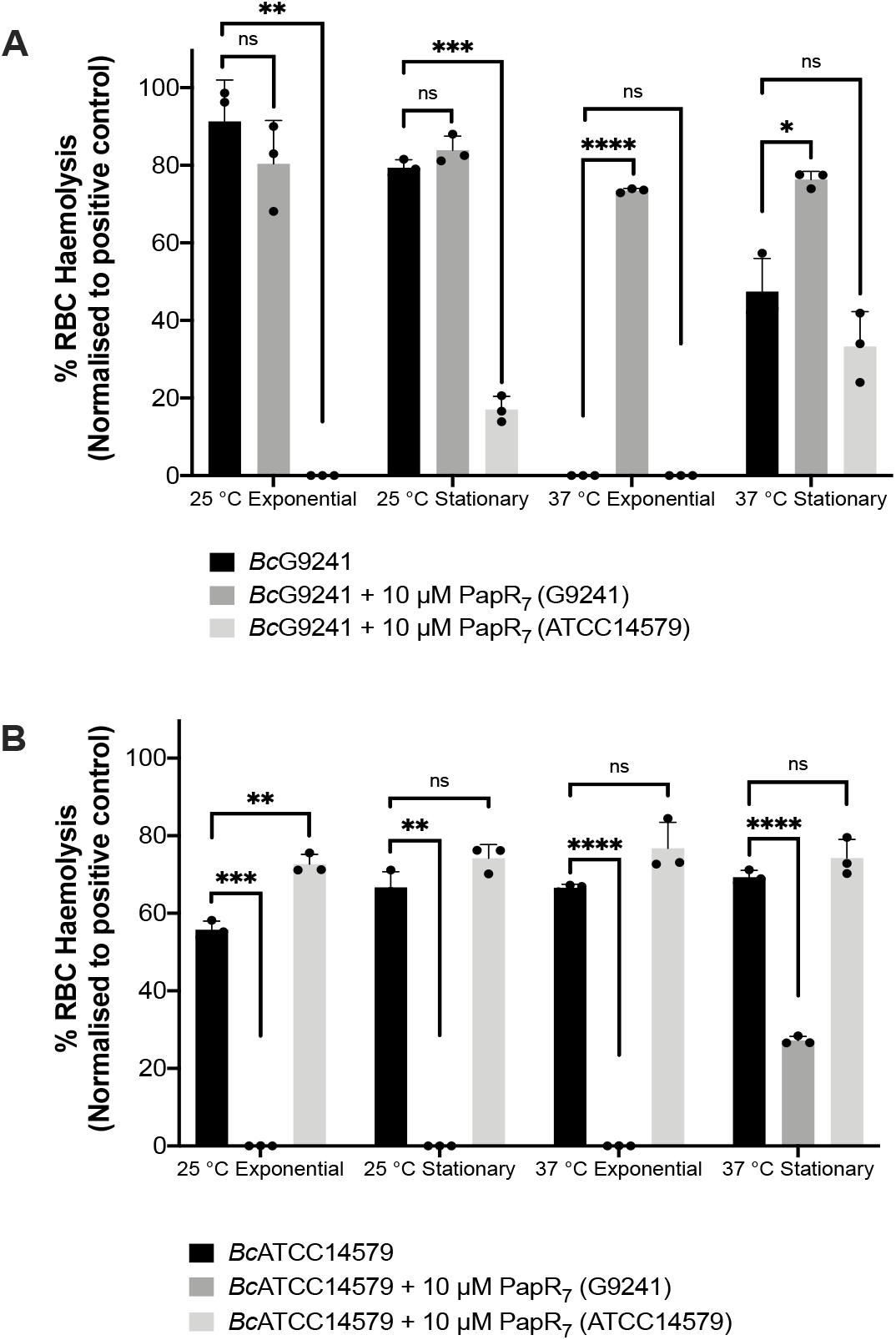
The effect of exogenous PapR_7_ in *Bc*G9241 and *Bc*ATCC14579. 10 *μ*M synthetic PapR_7_ (G9241 PapR_7_= SDLPFEH, ATCC14579 PapR_7_= KDLPFEY) were added when the bacterial culture was inoculated from OD_600_=0.005. Supernatant was extracted from mid-exponential and stationary phase growing *B. cereus* G9241. Supernatant was filter-sterilised and incubated with 4% RBCs for 1 hour. OD_540_ was measured and RBC lysis was calculated as a percentage of expected RBC lysis, normalised with 70 % lysis from 1 % (w/w) Triton X-100. Error bars denote one standard deviation, and all samples were to an n=3. * [P < 0.05], **[P < 0.01], ***[P < 0.001] and ****[P < 0.0001] as determined by unpaired t-test, with Welch’s correction.

Addition of exogenous non-self PapR_7_ molecules in *Bc*G9241 and *Bc*ATCC14579 (i.e., adding G9241 PapR_7_ into cultures of *Bc*ATCC14579 or adding ATCC14579 PapR_7_ into cultures of *Bc*G9241) led to a decrease in haemolytic activity at both temperatures (**Fig 4)**. This implies that the correct PapR_7_ is required for the expression of toxins, and that a non-self cognate variant of the peptide can actually interfere with the native PlcR-PapR circuit.

Subsequently, we wanted to observe how the addition of the synthetic PapR_7_ would affect the expression of PlcR-regulated toxins in *Bc*G9241 at 37 °C using the GFP reporters we have available. At 25 °C, addition of the cognate PapR_7_ to *Bc*G9241 led to a slight increase in expression of *nhe∷gfp*, no change in expression of *plc∷gfp* and *cytK∷gfp* and a decrease in expression of *hbl∷gfp*. A decrease in the expression of *hbl∷gfp* was still observed when lower concentrations of synthetic PapR were added to BcG9241 cultures (data not shown). At 37 °C, addition of cognate PapR_7_ to *Bc*G9241 led to a dramatic increase in expression of Nhe, Plc and CytK. This suggests that the processed PapR can indeed get imported into cells at 37 °C. There is no significant increase in *hbl∷gfp* expression with the addition of PapR_7_ at 37 °C (**Fig S12)** suggesting an additional level of regulation for these genes.

### The PapR maturation process is potentially preventing the expression of PlcR-controlled toxins in *Bc*G9241 at 37°C

The import of mature PapR does not appear to be a limiting factor involved in the temperature-dependent toxin expression phenotype. Consequently, there is a possibility that the protease(s) involved in processing PapR represents the limiting step within the PlcR-PapR circuit in *Bc*G9241. In *B. cereus*, it has been shown that the neutral protease NprB is involved in processing the pro-peptide PapR_48_ into the shorter and active form PapR_7_ (22). In the *B. cereus* and *B. thuringiensis* genome, the gene *nprB* is found adjacent to *plcR* and transcribed in the opposite orientation (6,21,22). To identify whether *Bc*G9241 and other *B. cereus-B. anthracis* “cross-over” strains have a functional copy of *nprB*, a synteny analysis using SyntTax (https://archaea.i2bc.paris-saclay.fr/SyntTax/Default.aspx) was carried out. SyntTax uses the genomic and taxonomic database obtained from NCBI. The NprB protein sequence from *Bc*ATCC14579 (RefSeq accession GCF_000007835.1) was used as the query protein. As shown in **Fig 5**, *Bc*G9241 as well as the “cross-over” strains *B. cereus* 03BB87, *B. cereus* 03BB102, *B. cereus* BC-AK and *B. cereus* bv *anthracis* CI can be seen to encode only remnants of the *nprB* gene located near the *plcR-papR* operon, with low synteny scores. In comparison, typical *B. cereus* sensu stricto and *B. thuringiensis* strains have intact copies of *nprB*, as previously described (6,21,22) with synteny scores above 97% (**Fig 5**). *B. anthracis* strains (Ames and Sterne) also lack the full copy of the *nprB* gene (22). This indicates that NprB may not be involved in processing PapR in *B. cereus* strains carrying functional copies of both *plcR* and *atxA*.

**Figure 5:**
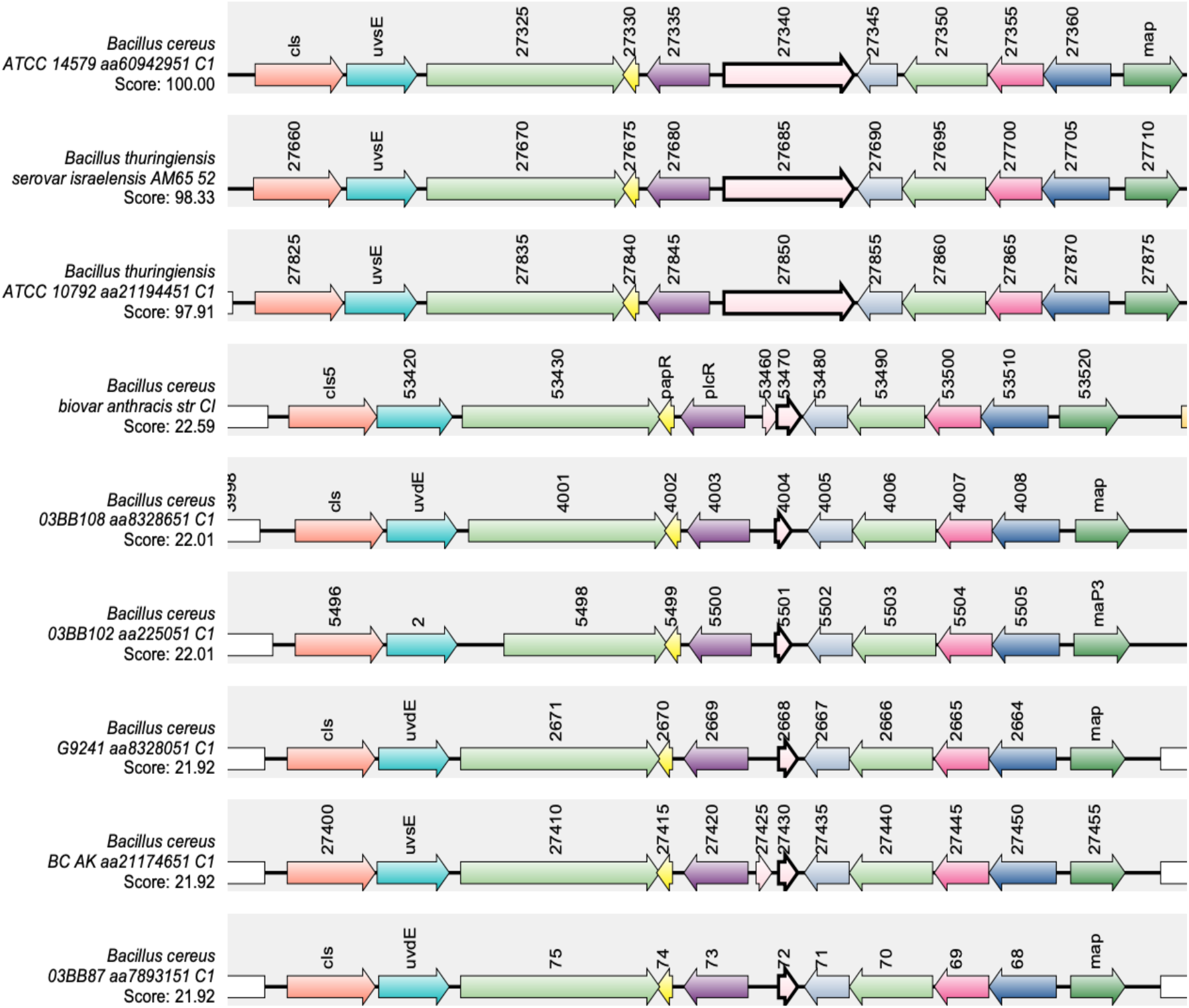
Synteny of the gene encoding *nprB* in *B. cereus* sensu stricto, *B. thuringiensis, B. weihenstephanensis, B. cereus* “cross-over” strains and *B. anthracis*. The NprB protein sequence from *Bc*ATCC14579 (RefSeq accession GCF_000007835.1) was used as the query protein. The gene encoding *nprB* is shown in pink with a bold border, *plcR* and *papR* are shown in purple and yellow, respectively. SynTax, a synteny web service, was used to look in the conservation of gene order (https://archaea.i2bc.paris-saclay.fr/SyntTax/Default.aspx).

This led us to question which protease(s) is/are capable of processing PapR in *Bc*G9241 and whether the temperature-dependent toxin expression in *Bc*G9241 is due to differential expression of these theoretical alternative protease enzymes. From the *Bc*G9241 secretome analysis of the supernatant extracted from cultures grown at 25 °C and 37 °C, several proteases were identified as highly expressed at 25 °C compared to 37 °C that could potentially be involved in processing PapR in *Bc*G9241 to its active form (**Table 1**). To determine whether temperature-dependent proteolytic activity is present in *Bc*G9241 as suggested by the secretome analysis, a protease activity assay was carried out using skim milk agar plates. Filtered supernatant of *Bc*G9241 grown at mid-exponential phase 25 °C showed hydrolysis of the skimmed milk casein whereas no clear zone was observed from cultures grown at mid-exponential phase 37 °C, which demonstrates that there is indeed a temperature-dependent protease activity deployed during mid-exponential phase of growth (**Fig 6**). Supernatants of *B. cereus* cultures into which synthetic PapR_7_ was added were also collected and spotted onto skim milk agar to look for any changes in proteolytic activity. Interestingly, the addition of the synthetic PapR_7_ peptide to cultures of either *Bc*G9241 or *Bc*G9241 ΔpBCX01 led to a significant increase in proteolytic activity at both temperatures (**Fig S13**), presumably caused by PlcR-regulated proteases such as the thermolysin metallopeptidase (AQ16_5317), which we have shown to be highly expressed at 25 °C compared to 37 °C (see above).

**Figure 6:**
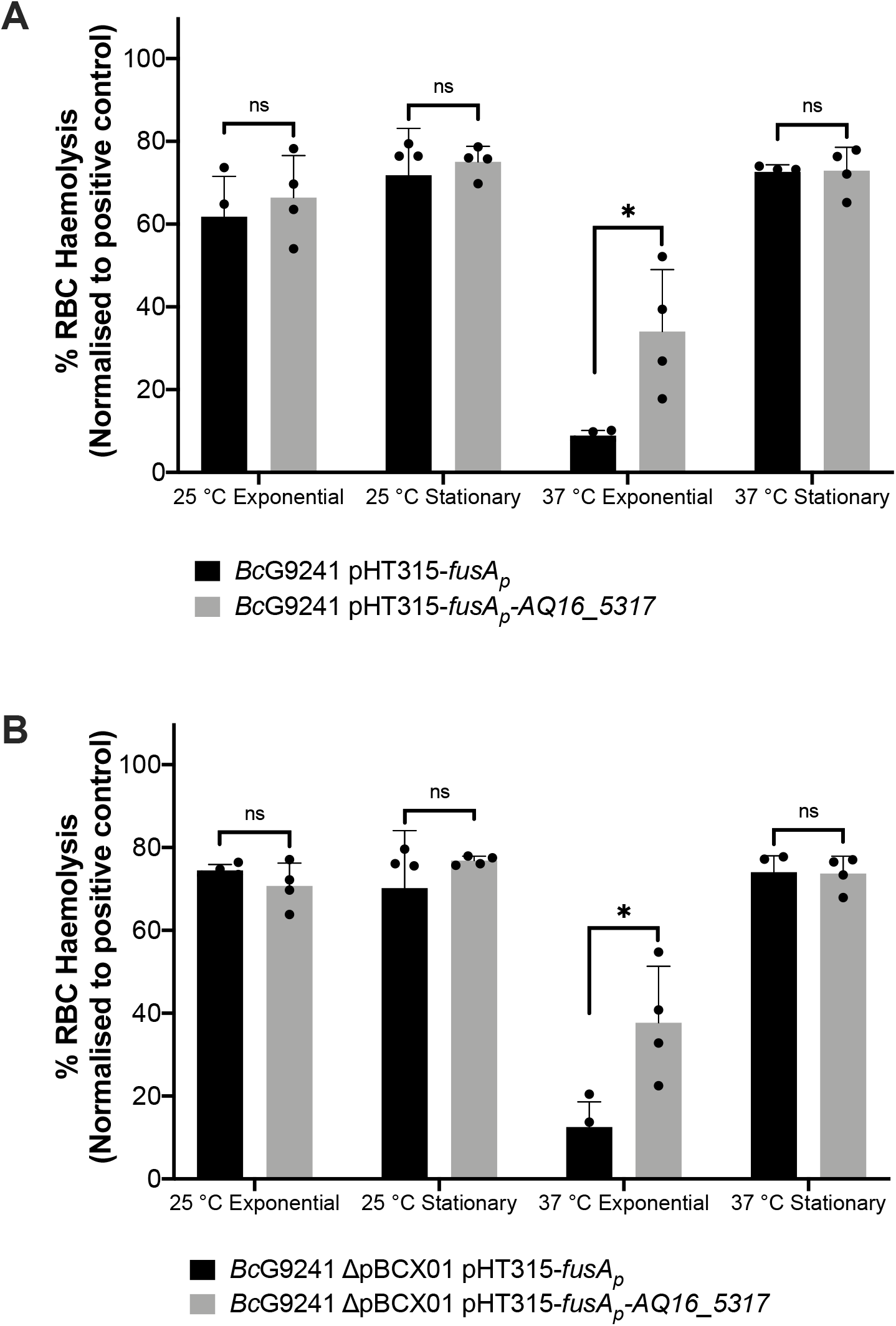
Slight increase in haemolytic activity with the presence of AQ16_5317 at 37 °C during exponential phase. The haemolysis assay was conducted by incubating the supernatant of (**A**) *Bc*G9241 and (**B**) *Bc*G9241 ΔpBCX01 with 4% RBC for one hour at 37 °C. OD_540_ was measured and RBC lysis was calculated as a percentage of expected RBC lysis, normalised with 70 % lysis from 1 % (w/w) Triton X-100. Stars above columns represent significance levels: * [P < 0.05] as determined by unpaired t-test, with Welch’s correction. Error bars denote one standard deviation, and all samples were to an n=4.

From the secretome analysis, AQ16_5317 labelled as a thermolysin metallopeptidase was found to be highly abundant at 25 °C compared to 37 °C during exponential and stationary phase (**Table 1**). Out of all these proteases/enzymes listed in **Table 1**, only the gene encoding the thermolysin metallopeptidase and collagenase has a PlcR-box on the promoter region. There is a possibility that AQ16_5317 is the protease involved in processing PapR in *Bc*G9241. From a synteny analysis to look into whether AQ16_5317 thermolysin metallopeptidase was present in other *B. cereus* species, a high synteny score as shown in some of the *B. cereus-B. anthracis* “cross-over” strains, as well as *B. cereus* sensu stricto, *Bacillus weihenstephanensis, B. anthracis* and *B. thuringiensis* (**Fig S14**).

To identify whether expressing AQ16_5317 at 37 °C would abolish the temperature-dependent haemolytic phenotype, the AQ16_5317 ORF with a promoter not linked to the PlcR-PapR regulator (*fusA* promoter) was cloned into the shuttle vector pHT315 which should be able to constitutively express the protease. When measuring the OD_600_ over 24 hours using a plate reader, strains containing the pHT315-*fusA_p_-AQ16_5317* construct did not alter the growth of the bacteria (data not shown). A haemolysis assay using sheep erythrocytes was carried out with *Bc*G9241 strains containing the constitutively expressed AQ16_5317. Cell free culture supernatants from cultures grown at 25 °C and 37 °C from mid-exponential and stationary phase were tested in this assay. Supernatants from 25 °C mid-exponential and early stationary phase cultures showed no change in haemolytic activity, with or without the constitutive expression of *AQ16_5317*. However, at 37 °C mid-exponential phase, constitutive expression of AQ16_5317 expressed at 37 °C led to a significant increase in haemolytic activity compared to the control (**Fig 6**). The implication being that this protease is indeed capable of processing PapR at 37 °C, leading to the increased expression of the PlcR regulon.

## DISCUSSION

The roles and expression of PlcR and AtxA are relatively well defined in *B. cereus* and *B. anthracis*, respectively. In *B. anthracis*, AtxA transcription and accumulation are enhanced at 37 °C compared to 28 °C (49). In *B. cereus* and *B. thuringiensis*, PlcR transcription has been observed at the onset of stationary phase, suggesting cell density is required for transcription of the regulon (6,50). Also, the *B. weihenstephanensis* KBAB4 is reported to exhibit temperature-dependent production of PlcR and PlcR-regulated toxins (51). However, due to the rare nature of some *B. cereus* strains containing both *plcR* and *atxA* (9,10,38–42), the understanding of how a bacterium such as *Bc*G9241 has incorporated two hypothetically conflicting virulence regulators (8) has not yet been studied in detail.

Haemolysis and cytolysis assays using the supernatant of *Bc*G9241 demonstrated lytic activity at 25 °C but not at 37 °C. The supernatant of *Bc*G9241 ΔpBCX01 also demonstrated temperature-dependent haemolytic and cytolytic activity, suggesting that this phenotype is not dependent on the pBCX01 virulence plasmid encoding AtxA1. In comparison, the supernatant of *Bt* Cry^−^ *ΔplcR* and *Ba St* showed little or no cytotoxicity against a variety of eukaryotic cells at both temperatures. As *Bt* Cry^−^ *ΔplcR* and *Ba St* lack a functional *plcR* gene, it supports the hypothesis that PlcR-regulated toxins are responsible. Together these findings led us to propose that *Bc*G9241 ‘switches’ its phenotype from a haemolytic *B. cereus*-like phenotype at 25 °C to a non-haemolytic *B. anthracis*-like phenotype at 37 °C.

The differential cytotoxicity pattern appears to be caused by the secretion of multiple cytolytic and haemolytic toxins at 25 °C, which includes Hbl, Nhe, Plc, CytK and a thermolysin metallopeptidase encoded by AQ16_5317, detected from the secretome analysis of exponentially grown *Bc*G9241 cells. All the corresponding genes encode an upstream PlcR box sequence (**Table S1**) and are known to be transcriptionally regulated by PlcR in *Bc*ATCC14579 (7,27). Using transcription-translation GFP reporters, we were able to confirm that *hbl∷gfp, nhe∷gfp, plc∷gfp* and *AQ16_5317∷gfp* are expressed in a temperature dependent manner, with higher expression at 25 °C in *Bc*G9241. This was also observed in *Bc*G9241 ΔpBCX01 (data not shown), further confirming that the temperature-dependent toxin production is independent of the virulence plasmid. In contrast, the expression of *hbl∷gfp, nhe∷gfp* and *BC_2735∷gfp* in *Bc*ATCC14579 were at a similar level between the two temperatures. Expression of *cytK∷gfp* in *Bc*G9241 was heterogeneous within the cell population, which Ceuppens *et al* (2012) also observed in *Bc*ATCC14579 using a cyan fluorescent protein reporter (52).

Interestingly, the most abundant proteins from the secretome analysis at 37 °C were phage proteins from the pBFH_1 phagemid. This is in agreement with the transcriptomic data carried out by our group (13),where high transcript levels of genes encoded on the pBFH_1 phagemid were identified at 37 °C compared to 25 °C from mid-exponentially grown *Bc*G9241 cells. It is possible that the expression of phage proteins might be interfering with normal PlcR-regulon toxin production. However, at this stage we have not confirmed whether phage protein expression is the cause or the effect of a loss of PlcR-mediated toxin expression at 37 °C, or indeed entirely independent.

The cell proteome analysis of mid-exponentially grown *Bc*G9241 cells revealed no accumulation of toxins at 37 °C, implying that temperature-dependent toxin expression is not regulated at the level of secretion. Also, PlcR was detected at both temperatures from the cell proteome analysis with no significant difference between expression levels. Consequently, it can be concluded that the temperature-dependent toxin profile is not due to levels of PlcR in the cell. Instead, this suggests that the control point for temperature-dependent toxin secretion could be due to differential activity of the PlcR-PapR active complex.

In *Bc*G9241 and *Bc*ATCC14579, expression of PlcR using transcription-translation GFP reporters was first observed at the onset of stationary phase, in agreement with previous observations in *B. thuringiensis* (6). Expression of PlcR was highly heterogeneous during the onset of stationary phase and by 24 hours. As PlcR is under the direct- and indirect influence of other transcriptional regulators such as Spo0A and CodY (27,53), it is possible that these regulators play a role in the heterogeneous expression of PlcR. Phosphorylated Spo0A is able to inhibit the expression of PlcR due to the presence of two Spo0A-boxes between the PlcR box in the promoter region of *plcR* (27), while CodY controls the Opp system involved in importing processed PapR into the cell to activate PlcR (53). Population heterogeneity between genetically identical cells could be beneficial in order to survive, persist in fluctuating environment or be helpful for division of labour between cells. Transcription-translation expression of PapR using GFP reporters was also observed across the growth phase. Unexpectedly, expression of PapR in *Bc*G9241 increased dramatically at 37 °C, with a near homogenous expression observed by 24 hours. This is in contrast with what is observed in *Bc*ATCC14579, where the expression of PapR was heterogeneous at 25 °C and 37 °C. It is possible that at 37 °C, *Bc*G9241 cells are trying to compensate the low expression of the PlcR regulon by expressing PapR highly.

As analysis of the cell proteome did not show a build-up of toxins at 37 °C, we wanted to identify whether the temperature-dependent toxin production was caused by a limiting step within the PlcR-PapR regulatory circuit in *Bc*G9241: import of mature PapR or processing of immature PapR.

Addition of synthetic PapR_7_ to *Bc*G9241, which would bypass the secretion and processing of the full-length peptide restored haemolytic activity at 37 °C. Supplementing the non-cognate form of the PapR_7_ peptide into *B. cereus* strains led to suppression of haemolytic activity at both temperatures, confirming that the activating mechanism of PlcR-PapR is strain specific. This observation has been previously noted in *B. thuringiensis* (28). Using the PlcR-regulated toxin reporter strains made in this study, the addition of synthetic PapR_7_ led to an increase in *nhe∷gfp, plc∷gfp* and *cytK∷gfp* expression at 37 °C in *Bc*G9241. This further confirms that the mature form of PapR can be imported into the cell at 37 °C in order to bind to PlcR and express the PlcR regulon. There was no significant increase in *hbl∷gfp* expression at 37 °C with the addition of PapR_7_. This suggests that Nhe, Plc and CytK are responsible for the haemolytic activity of *Bc*G9241 observed at 37 °C in the presence of PapR_7_ (**Fig 4**). Intriguingly, the addition of PapR_7_ at 25 °C led to a decrease in the expression of *hbl∷gfp* in *Bc*G9241. There is a possibility that there are other regulators that play a role in the expression of this enterotoxin such as ResDE (redox regulator), FnR, RpoN and Rex (54,55). It has been demonstrated that FnR, ResD and PlcR are able to form a ternary complex *in vivo* (56), which could explain the decrease of Hbl expression when synthetic PapR_7_ were added.

Finally, this led us to question as to whether the processing of PapR by an extracellular protease(s) was the limiting step causing the temperature-dependent toxin expression. The gene *nprB*, which encodes for the neutral protease involved in processing PapR in *B. cereus* and *B. thuringiensis* is truncated in *B. anthracis* (22), as well as in *Bc*G9241 and some of the *B. cereus-B. anthracis* “cross-over” strains that carry functional copies of *plcR-papR* and *atxA* (9,10,38–42). The loss of a functional copy of *nprB* may have contributed to the accommodation of *atxA* in these strains and potentially allowed *B. cereus-B. anthracis* “cross-over” strains to carry both regulators. Temperature-dependent proteolytic activity was observed using the supernatant of *Bc*G9241, suggesting that the processing of PapR by extracellular proteases may be temperature-dependent manner. From the secretome analysis of *Bc*G9241, AQ16_5317, a thermolysin metallopeptidase, was found to be one of the most highly expressed proteases at 25 °C compared to 37 °C. Constitutive expression of AQ16_5317 led to a slight increase in haemolytic activity at 37 °C, suggesting that AQ16_5317 is capable of processing PapR into its mature form leading to the expression of PlcR-regulated toxins. The reason for not observing a similar level of haemolytic activity as observed using the supernatant extracted from 25 °C growth culture could be that AQ16_5317 may require further processing in order to be in its active form or is not stable enough to carry out its function at 37 °C. It is also likely that though AQ16_5317 is able to process PapR and express PlcR-regulated toxins at 37 °C, other proteases that have not been studied here can also carry out this function, and therefore further analysis is required. The possibility of other proteases processing immature PapR has been stated by Slamti *et al* (2014), though data have not been published to support this statement (57).

Overall, this study reveals that haemolytic and cytolytic activity in *Bc*G9241 is determined by temperature. Lytic activity at 25 °C was accompanied by higher levels of PlcR-regulated proteins including Hbl, Nhe, Plc, CytK and AQ16_5317, a thermolysin metallopeptidase. Production of these virulence factors at 25 °C may be essential for the invasion of insect hosts. As shown in **Figure 7**, another finding of our work is that the temperature-dependent toxin production is due to differential expression of protease(s) involved in processing the immature PapR into its mature form to be reimported and and then activate PlcR. This study suggests that temperature-dependent regulation of the PlcR-PapR regulator allows *Bc*G9241 to accommodate a functional copy of *atxA*. We hypothesise that this has led to the ability of *Bc*G9241 to switch between a *B. cereus*-like phenotype at 25 °C and a *B. anthracis*-like phenotype at 37 °C. The lower activity of the PlcR regulon at 37 °C compared to 25 °C could be to allow the expression of AtxA and its regulon, known to be expressed at 37 °C in *B. anthracis* (49). The characterisation of this “cross-over” strain demonstrates that the evolution of *B. anthracis* as a significant mammalian pathogen is not merely about acquisition of genetic information but is also a story of the power of regulation in controlling potential incompatibilities between incumbent and newly acquired systems which may be a feature of other emerging pathogens.

**Figure 7:**
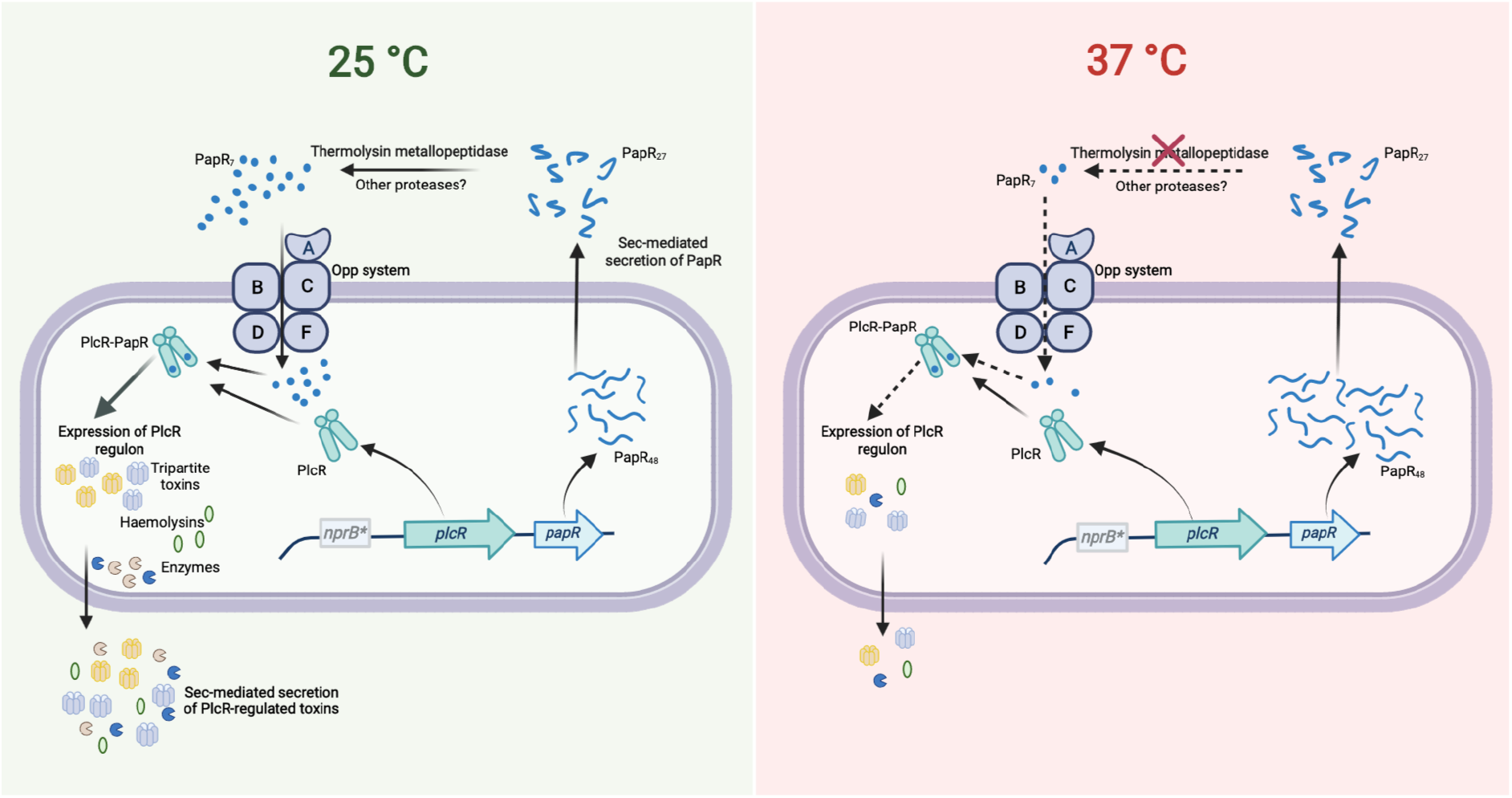
The PlcR-PapR regulation circuit in *Bc*G9241 at 25 °C and 37 °C. In *Bc*G9241, the expression of *plcR* was observed at 25 °C and 37 °C, suggesting that rather than the expression of the regulator, the activity of the PlcR-PapR active complex is causing differential production of PlcR-regulated toxins. PapR was highly expressed at 37 °C compared to 25 °C, potentially to compensate for the low expression of the PlcR regulon. Secretion of PapR was observed at both temperatures, suggesting that the Sec machinery is functional at both temperatures. The import system is also functional at both temperatures when PapR_7_ is available extracellularly. Remnants of the *nprB* gene (*nprB*\*) are present in *Bc*G9241, thus the protease NprB is not involved in the maturation of PapR as observed in *B. cereus* and *B. thuringiensis*. A PlcR-regulated thermolysin metallopeptidase (AQ16_5317) was identified to have the ability to process PapR and cause haemolytic activity. AQ16_5317 is highly expressed at 25 °C compared to 37 °C. Diagram created with BioRender.com

## MATERIALS AND METHODS

### Bacterial strains and growth conditions

Bacterial strains used in this study were *Bc*G9241 (9), *Bc*G9241 ΔpBCX01 (13), *B. cereus* reference strain ATCC 14579 (American Type Culture Collection, Manassas, Va.), the *plcR*-defective strain *B. thuringiensis* 407 Cry^−^ Δ*plcR* (18), and *B. anthracis* Sterne 34F2 (pXO1^+^, pXO2^−^). *Bacillus* strains were cultured in 5 mL lysogeny broth (LB) at either 25 °C or 37 °C overnight before subculturing into 5 mL LB media. All cultures were incubated with shaking at 200 rpm. Larger cultures of *Bacillus* strains were cultured in 50 mL of LB unless otherwise specified. For cloning, *Escherichia coli* DH5-*α* (NEB) and the methylation deficient *E. coli* ET12567/pUZ8002 (58) were used in this study. *E. coli* strains were grown in 5 mL LB media at 37 °C, shaking at 200 rpm. Media contained antibiotics when appropriate: ampicillin (100 μg/mL), chloramphenicol (25 μg/mL), kanamycin (25 μg/mL) for *E. coli* and erythromycin (25 μg/mL) for *B. cereus* strains.

### Haemolysis assay

Haemolytic activity was determined from sheep erythrocytes as described in (60). Briefly, erythrocytes were diluted to 4% (vol/vol), in RPMI-1640 medium and 50 μL of this cell suspension were transferred to a 96-well round-bottom polystyrene plate and incubated with 50 μL of filtered supernatants of *B. cereus* cells grown to exponential (OD_600_=0.5) or stationary phase (OD_600_=0.5). Following a 1 h-incubation at 37 °C, lysis of human/sheep erythrocytes were determined by quantifying the haemoglobin release by measurement of the absorbance at 540 nm in the resulting supernatant. LB and 1% Triton X-100 were used as negative and positive control for 0% lysis and 70% lysis, respectively. %RBC haemolysis was calculated as (OD_sample_ − OD_negative control_)/(OD_positive control_ − OD_negative control_) × 70%. Assays were done by triplicate unless otherwise stated.

### Protein extraction

Before cultures were seeded for protein extraction, pre-cultures of *Bc*G9241 were used to synchronise bacterial cell growth. Pre-cultures were inoculated into 50 ml of LB broth at OD_600_ = 0.005, for protein extraction. Secreted proteins were collected from mid-exponential phase or late stationary phase at both 25 °C and 37 °C. Once *Bc*G9241 had grown to the appropriate time point, 6.75 OD units of cells were centrifuged for 5 minutes at 8000 rpm at 4 °C.

#### (i) Protein extraction for secretome proteomics using in-gel digestion

Supernatant was extracted and acidified to pH 5 using 10% trifluoric acid (TFA). 50 μl of StrataClean resin (Agilent) was added to each sample before vortexing for 1 minute. All samples were incubated overnight on a rotor wheel mixer overnight at 4 °C for efficient protein extraction. StrataClean resin was collected by centrifugation at 870 g for 1 minute. Cell supernatant was removed, and the beads resuspended in 100 μl of Laemlli buffer. The suspension was boiled at 95 °C for 5 minutes, to unbind the protein from the resin. Beads were pelleted at 870 g for 1 minute and protein-Laemlli buffer suspension collected.

25 μl of the secreted proteins were ran on a Mini-PROTEAN® TGX™ precast gel (Bio-Rad). The whole lane of the gel for each sample was sliced into 4 mm sections and washed with 1 ml of 50% ethanol in 50 mM ammonium bicarbonate (ABC). This wash was incubated for 20 minutes at 55 °C, shaking at 650 rpm. The wash solution was removed and this step was repeated twice more. The gel was dehydrated in 400 μl of 100% ethanol by incubation at 55 °C for 5 minutes, with 650 rpm shaking. Once the gel was dehydrated, remaining ethanol was removed. Disulphide bonds were reduced by addition of 300 μl of 10 mM dithiothreitol (DTT) in 50 mM ABC. This was incubated for 45 minutes at 56 °C with 650 rpm shaking. DTT was removed and samples were cooled to room temperature. Cysteine residues were alkylated by adding 300 μl of 55 mM iodoacetamide (IAA) in 50 mM ABC with incubation at room temperature, in the dark for 30 minutes. IAA was removed and gel was washed as before by adding 1 ml of 50% ethanol in 50 mM and incubated at 55 °C for 20 minutes with shaking at 650 rpm. The ethanol was removed and this wash was repeated twice. Gel pieces were again dehydrated with 400 μl of 100% ethanol and incubated for 5 minutes at 55 °C. 200 μl of trypsin at 2.5 ngμl^−1^ was added to the dehydrated gel and ABC added to ensure the rehydrated gel was fully submerged. The trypsin digest was incubated for 16 hours at 37 °C with 650 rpm shaking. The digest was stopped by addition of 200 μl 5% formic acid in 25% acetonitrile. The solution was sonicated for 10 minutes at 35 KHz and the supernatant extracted. This step was repeated three more times. A C18 stage-tip (Thermo Scientific™) was made and conditioned by centrifuging 50 μl 100% methanol through the tip for 2 minutes at 2000 rpm. 100% acetonitrile was washed through the tip in the same manner to equilibrate it. The tip was further equilibrated with 2% acetonitrile with 1% TFA washed through the tip as before but for 4 minutes. Samples were then diluted to a concentration of 10 μg of protein in 150 μl final volume of 2% acetonitrile/0.1% TFA. Samples were collected on the stage tip by centrifugation through the stage tip for 10 minutes under previous spin conditions. The membrane was washed with 50 μl 2% acetonitrile/0.1% TFA by centrifugation at 2000 rpm for 4 minutes. Peptides were eluted in 20 μl 80% acetonitrile. Samples were dried to a total volume of 40 μl at 40 °C in a speed-vac. Samples were resuspended in 55 μl of 2.5% acetonitrile containing 0.05% TFA and sonicated for 30 minutes at 35 KHz. Samples were dried to a total volume of 40 μl at 40 °C in a speed-vac again ready for mass spectroscopy. Nano liquid chromatography-electrospray ionisation-mass spectrometry (nanoLC-ESI-MS)/mass spectrometry (MS) was used to carry out the analysis.

#### (ii) Protein extraction for intracellular proteomics using in-urea protein digests

Cell supernatant was removed and cell pellets were suspended in 100 μl of 8M urea. Suspensions were transferred to Lysing Matrix B tubes (MP Biomedicals) and cells were lysed using the FastPrep®-24 Classic instrument with a COOLPREP™ adapter (MP Biomedicals). Bead beating was conducted at 6 ms^−1^ for 40 s for 2 cycles, with a 300 s pause between cycles. Samples were filtered through nitrocellulose membranes to remove the beads and protein was quantified using a Qubit 2.0 fluorometer and a Qubit™ protein assay kit (Life Technologies). 50 μg of protein sample was suspended in 50 μl of 8 M urea buffer. 5.5 μl of 10 mM DTT was added the samples were incubated for 1 hour at room temperature. 6.2 μl of 55 mM IAA was added to samples before 45 minutes incubation at room temperature in the dark. Samples were then diluted to 100 μL total volume by addition of 50 mM ABC. 1 μg of trypsin was added to each sample per 50 μg protein and incubated for 16 hours at room temperature. Samples were filtered through a C-18 stage tip as described previously and concentrated to 40 μl in a speed-vac, ready for mass spectroscopy. nanoLC-ESI-MS/MS was used to carry out the analysis.

### Perseus analysis of proteomics data

The Perseus software platform (Max Planck Institute) was used to analyse the highly multivariate proteomics data. Peptides only identified by site, reversed peptide sequences and potential contaminants were filtered out. Secretome data was normalised by the mean label-free quantification (LFQ) intensity value. Whole cell proteomics data was normalised by median as the data was normally distributed. Protein hits were filtered out if they didn’t have 3 values in at least one condition measured. Volcano plots were plotted using a p value = 0.05 and a log2-fold change = 1.

### Generation of plasmid-based transcription-translation GFP reporters

Constructs made for this study are listed on **Table S2**. Transcription-translation fusions with the *gfp* gene were constructed by PCR in a pHT315 vector (59) containing *gfp* (pHT315-*gfp*). The vector was linearized using appropriate restriction enzymes (NEB) and purified after agarose gel electrophoresis using the GFX™ PCR DNA and Gel Band Purification Kit (GE Healthcare). Insert fragments were amplified with Q5 DNA polymerase (NEB) by PCR with the appropriate primer pairs listed on **Table S3**. The resulting fragments were digested with the appropriate restriction enzymes, purified after agarose gel electrophoresis using the GFX™ PCR DNA and Gel Band Purification Kit (GE Healthcare) and ligated into the linearized pHT315-*gfp* vector. Plasmid constructs were transformed into chemically competent *E. coli* DH5-α cells through heat shock. Once confirmed by DNA sequencing, all vectors were transformed into the non-methylating *E. coli* ET12567 strain by electroporation (**Supplementary Materials and Methods**). Vectors amplified by *E. coli* ET12567 were purified and transformed into *B. cereus* strains using electroporation (**Supplementary Materials and Methods**).

### Fluorescent reporter strain assays

For growth curves and fluorescence measurements, *B. cereus* strains were sub-cultured at a starting OD_600_ of 0.05 into a clear flat bottom 96-well plate (Greiner) containing 100 μL of LB media per well. Cultures were grown in a FLUOstar Omega microplate reader (BMG LabTech) at either 25 °C or 37 °C with continuous orbital shaking at 700 rpm. Absorbance measurements (OD_600_) and fluorescence intensity (excitation filter = 482 nm and emission filter = 520 nm for GFP) were taken hourly for 24 hours. Each plate contained *Bc*G9241 and *Bc*ATCC14579 strains carrying GFP reporters as well as each strain carrying a control plasmid with no promoter upstream of *gfp* (pHT315-*gfp*). The fluorescence of all readings was first normalized to the fluorescence of blank media samples and then normalized by subtracting the autofluorescence of the corresponding control strain. The rate of change in fluorescence (ΔGFP/OD_600_) using the data obtained from the microplate reader was calculated by subtracting the fluorescence at a given time point by the fluorescence of the previous time point:

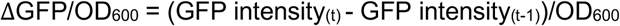

### Peptide Synthesis

Peptides SDLPFEH (G9241 PapR_7_) and KDLPFEY (ATCC14579 PapR_7_) were synthesised by GenScript (USA) at a purity >98% and diluted with sterile nuclease-free water. All experiments with the use of PapR_7_ were added at a concentration of 10 μM and during lag growth phase (OD_600_ = 0.1), unless otherwise stated.

### Light and Fluorescence Microscopy

1 % agarose in water were made and heated using a microwave until the agarose has completely dissolved. 200 *μ*l of molten agarose was added onto a microscope glass slide and a coverslip placed on top. When the agarose pad has dried and the sample is ready for observation, 2 μl of sample was applied to a prepared agarose pad and a cover slip placed over them. Images were captured on a Leica DMi8 premium-class modular research microscope with a Leica EL6000 external light source (Leica Microsystems), using an ORCA-Flash4.0 V2 Digital CMOS Camera (Hamamatsu) at 100x magnification.

## PapR_7_ activity assay using PlcR-regulated toxin reporters

*Bc*G9241 and *Bc*ATCC14579 containing PlcR-regulated toxin GFP reporters were grown overnight in LB medium with selective antibiotics. Mid-exponentially grown pre-cultures of *B. cereus* strains containing PlcR-regulated toxin reporters were diluted to OD_600_ 0.01 and 10 μM of PapR_7_ were added. In a black tissue culture treated 96-well microtiter plate (Greiner, Scientific Laboratory Supplies), 100 μL of the culture were added in each well and the GFP intensity and OD_600_ were measured every hour for over 24 hours using the Omega FluoSTAR (BMG LabTech) microplate reader.

## AUTHORS AND CONTRIBUTORS

**Shathviga Manoharan^1^**: Planned and performed experiments and wrote much of the manuscript

**Grace Taylor-Joyce^1^**: Assisted in experiments and wrote parts of the manuscript

**Thomas Brooker^1^**: Planned and performed some of the experiments.

**Carmen Sara Hernandez-Rodrıguez^2^**: Planned and performed some of the experiments.

**Les Baillie^3^**: Provided certain bacterial strains and provided advice on handling them.

**Petra Oyston and Victoria Baldwin^4^**: Provided advice on handling the pathogenic strains and assisted in interpreting the results.

**Alexia Hapeshi^1^**: Assisted in some experimental work and in interpreting certain results.

**Nicholas R. Waterfield^1^**: Experimental planning, secured funding, assisted in interpreting results and provided guidance and edits for writing the manuscript.

^1^Division of Biomedical Sciences, Warwick Medical School, University of Warwick, Gibbet Hill Road, Coventry, CV4 7AL, United Kingdom

^2^Institut Universitari de Biotecnologia i Biomedicina, Departament de Genètica, Facultad de Ciències Biològiques, University of Valencia, 46100 Burjassot, Valencia, Spain

^3^School of Pharmacy and Pharmaceutical Sciences, Cardiff University, CF10 3AT, Cardiff, United Kingdom

^4^CBR Division, Dstl Porton Down, Salisbury, SP4 0JQ, United Kingdom

## ACKNOWLEDGEMENTS

We would like to thank the Dstl for providing funding and guidance throughout the project including valuable quarterly update meetings. We would also like to acknowledge the contribution of the WPH Proteomics Research Technology Platform (Gibbet Hill Road, University of Warwick, UK).

