## Supplementary figures and tables for "From *cereus* to anthrax and back again: The role of the PlcR regulator in the “cross-over” strain *Bacillus cereus* G9241" (from another version of this preprint: version 1, 2022-07-07)

**Supplementary Data**

Shathviga Manoharan^1^, Thomas Brooker^1^, Carmen Sara Hernandez-Rodriguez^2^, Grace Taylor-Joyce^1^, Alexia Hapeshi^1^, Victoria Baldwin^3^, Les Baillie^4^, Petra C. F. Oyston^3^ and Nicholas R. Waterfield^1^**^♦^**.

^1^Division of Biomedical Sciences, Warwick Medical School, University of Warwick, Gibbet Hill Road, Coventry, CV4 7AL, United Kingdom

^2^Dpto. Microbiología y Ecología, Instituto BIOTECMED, Universitat de València, 46100 Burjassot, Spain.

^3^CBR Division, Dstl Porton Down, Salisbury, SP4 0JQ, United Kingdom

^4^School of Pharmacy and Pharmaceutical Sciences, Cardiff University, CF10 3AT, Cardiff, United Kingdom

**Supplementary Figures**

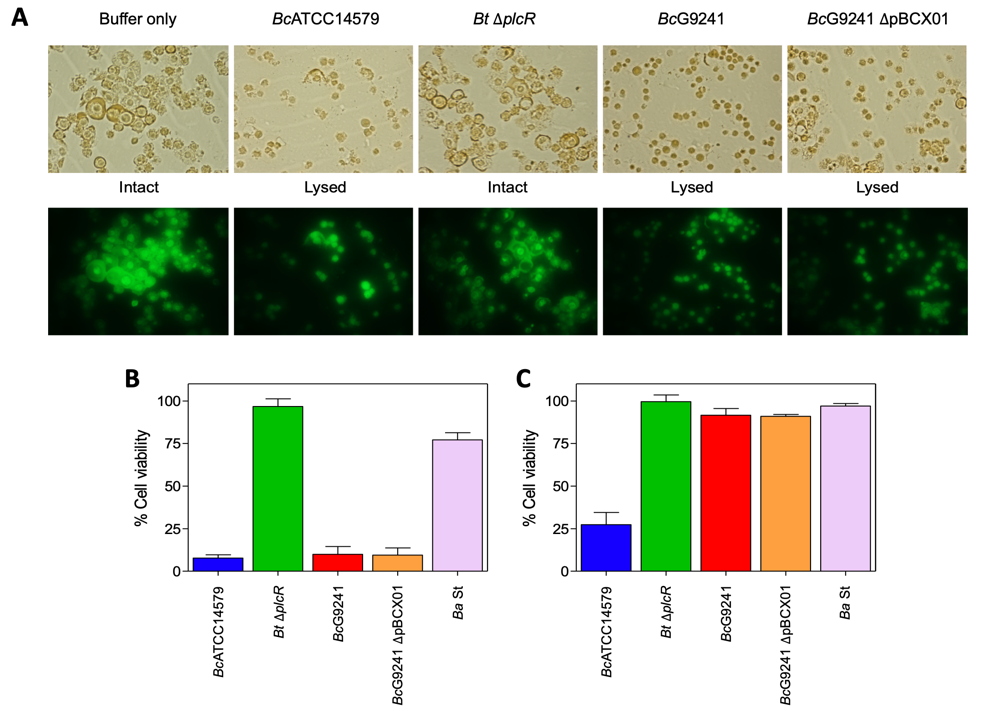
**Supplementary Figure S1: *Bc*G9241 supernatant is more toxic to *ex vivo M. sexta* haemocytes, when extracted from 25 °C growing culture compared to 37 °C growing culture.** (**A**) Micrographs of *ex vivo* *M. sexta* haemocytes with culture supernatants of *Bc*ATCC14579, *Bt* $\Delta$*plcR*, *Bc*G9241 and *Bc*G9241 $\Delta$pBCX01 added at 25 ºC. Cells were incubated with acridine orange (10 μg/mL) for two minutes and then viewed and imaged with filters for green fluorescence in a laser-scanning confocal microscope (Zeiss)**.** To quantify the effect of supernatants from these same strains using haemocyte cell viability assays, bacterial strains were cultured for 16 h in liquid LB media at (**B**) 25 ºC and (**C**) 37 ºC before testing on freshly prepared *ex vivo* *M. sexta* haemocytes. *Bc*ATCC14579 refers to *B. cereus* ATCC14579*, Bt* $\Delta$*plcR* refers to *B. thuringiensis* 407 Cry^-^ $\Delta$*plcR, Bc*G9241 refers to *B. cereus* G9241, *Bc*G9241 $\Delta$pBCX01 refers to *B. cereus* G9241 $\Delta$pBCX01 and *Ba* St refers to *B. anthracis* Sterne. Error bars denote standard deviation, and all samples were to an n=3.

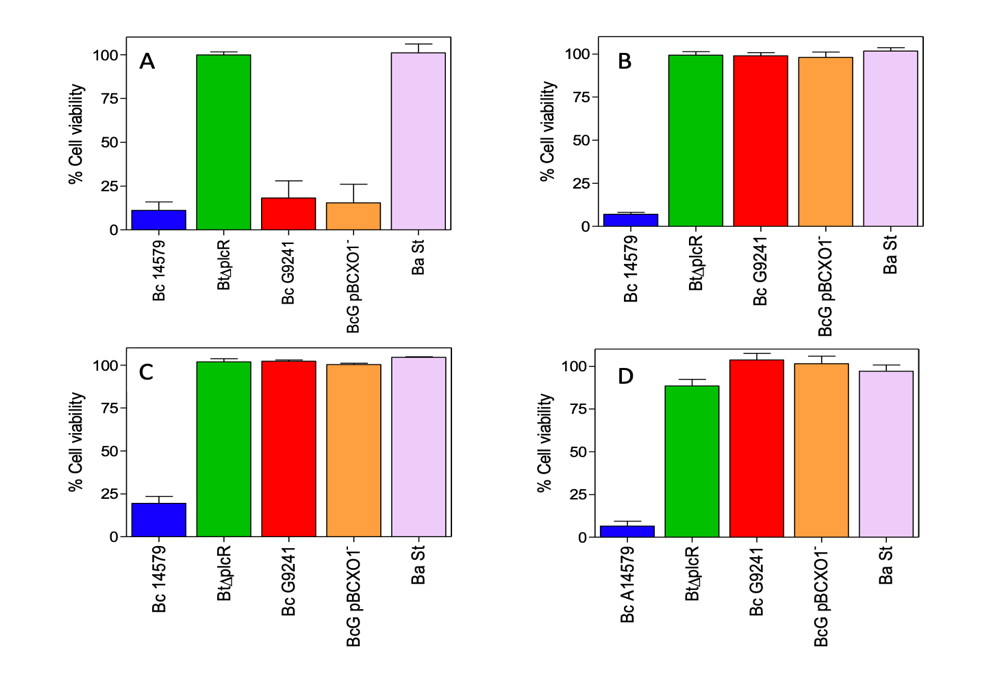

**Supplementary Figure S2: *Bc*G9241 supernatant is more toxic against a range of eukaryotic cells when extracted from 25 °C growing culture compared to 37 °C growing culture.** % Cell viability of T2-lymphocytes tested against neat supernatants of *Bacillus* cultures grown for 16 h at (**A**) 25 ºC or (**B**) 37 ºC. The effect of neat supernatants of *Bacillus* cultures grown at 37 ºC were also tested on (**C**) polymorphonuclear leukocytes (PMNs) and (**D**) THP-1 activated macrophages. Note the cytotoxicity pattern of PMNs and macrophages treated with 25ºC grown supernatants, showed the same trend as (**A**) and so is omitted for brevity. Strain name key is similar to that shown in Supplementary Figure S1. Error bars denote standard deviation, and all samples were to an n=3.

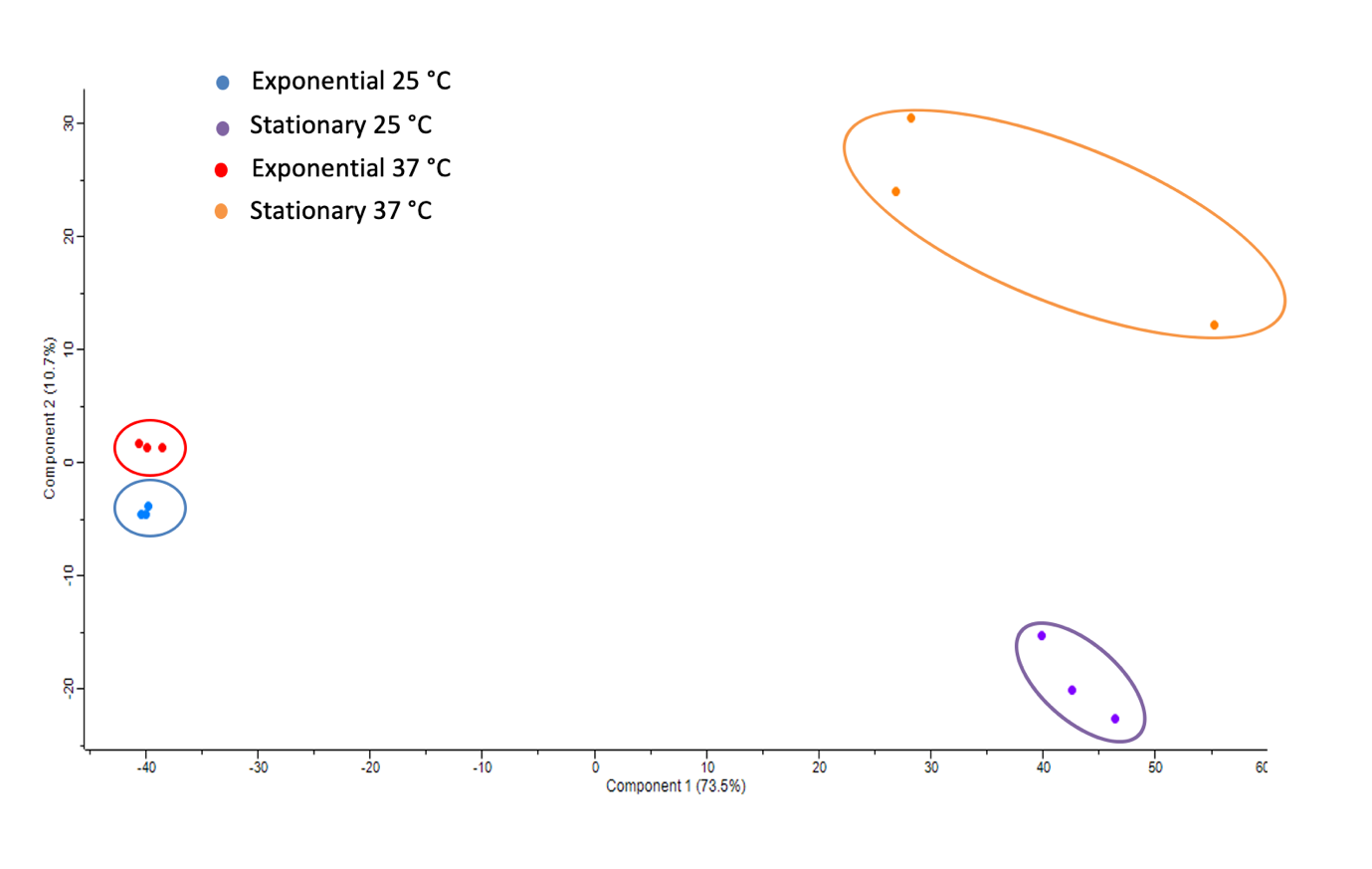

**Supplementary Figure S3: A principal component analysis of secretome of *Bc*G9241.** Proteins were extracted from the supernatant of *Bc*G9241 growing exponentially and in stationary phase at both 25 °C and 37 °C. The PCA plot was produced using Perseus software (Max Planck Institute). Three dots of each colour represent the biological replicates under each condition.

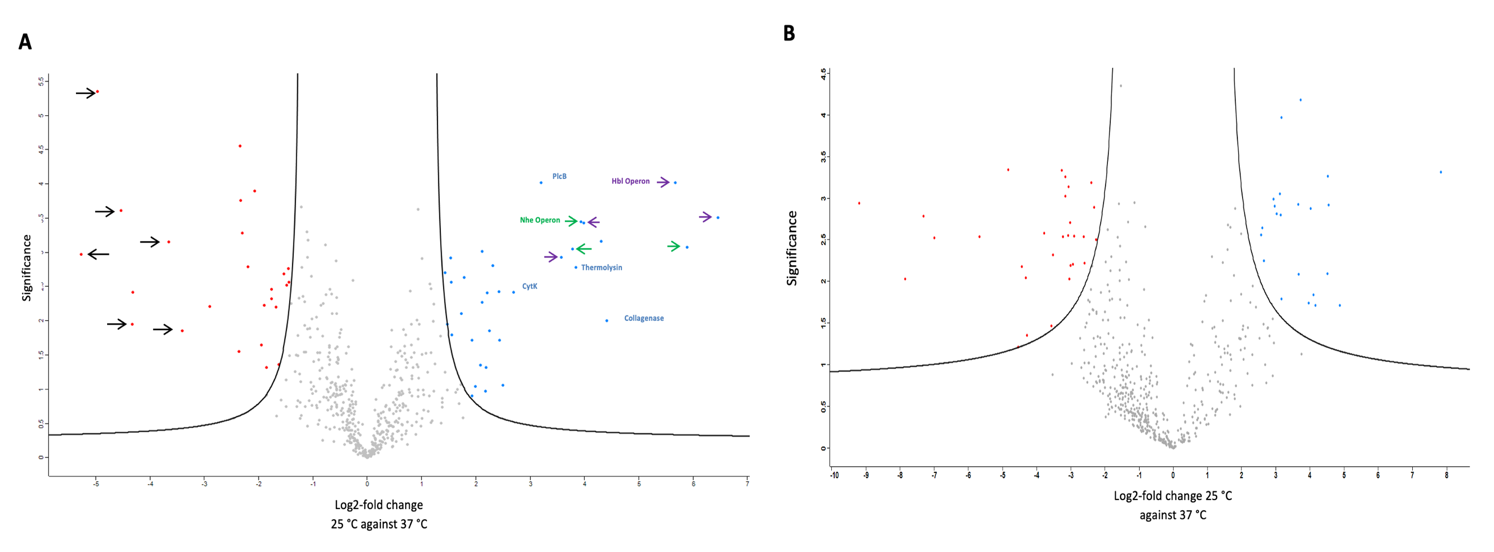

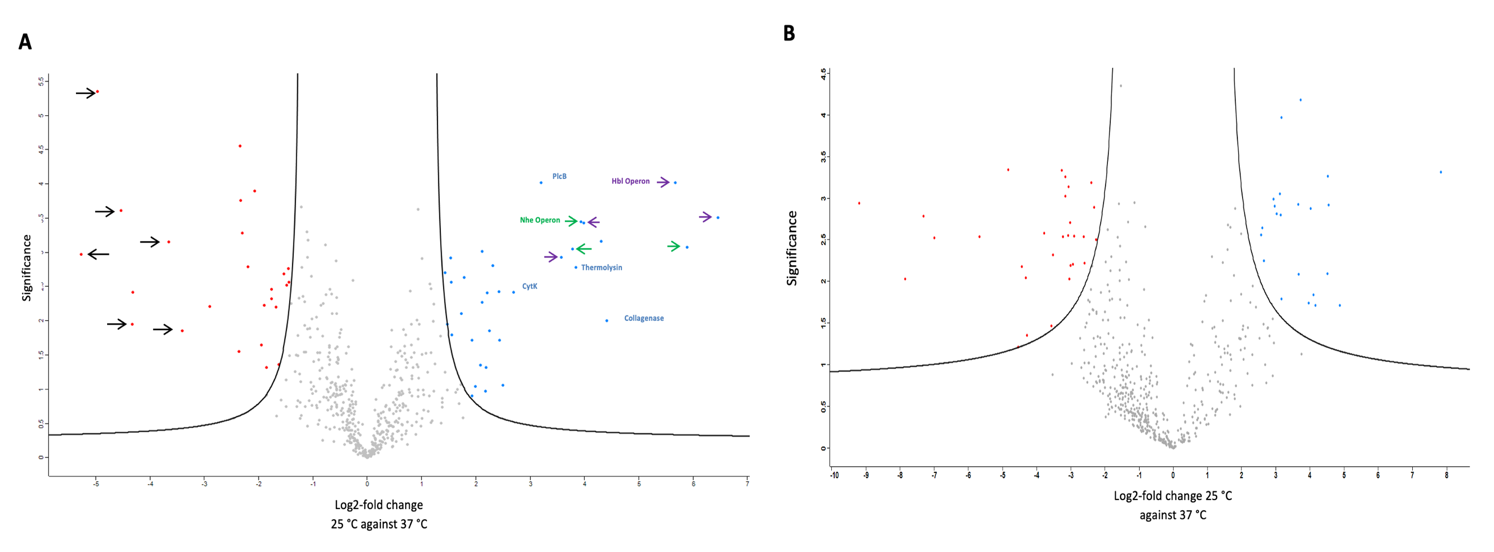
**Supplementary Figure S4: (A) A volcano-plot comparison of *Bc*G9241 mid-exponential secretome when grown at 25 °C and 37 °C.** *Bc*G9241 secretes a cocktail of toxins at 25 °C compared to 37 °C when growing exponentially. All samples were collected at n=3. Red and blue dots show proteins more highly secreted at 37 °C and 25 °C respectively. The black curved lines represent the significance cut off criteria of a P-value of <0.05 and a 2-fold change in protein level. **(B) A volcano-plot comparison of *Bc*G9241 stationary phase secretome when grown at 25 °C and 37 °C.** Mass spectroscopy was used to analyse the secreted proteins of *Bc*G9241 WT, in stationary phase at both 25 °C and 37 °C in LB broth, 200 rpm. Stationary phase growth occurred after 7 and 10 hours growth at 37 °C and 25 °C respectively. All samples were collected at n=3. Red and blue dots show proteins more highly secreted at 37 °C and 25 °C respectively. The black curved lines represent the significance cut off criteria of a p-value of <0.05 and a 2-fold change in protein level.

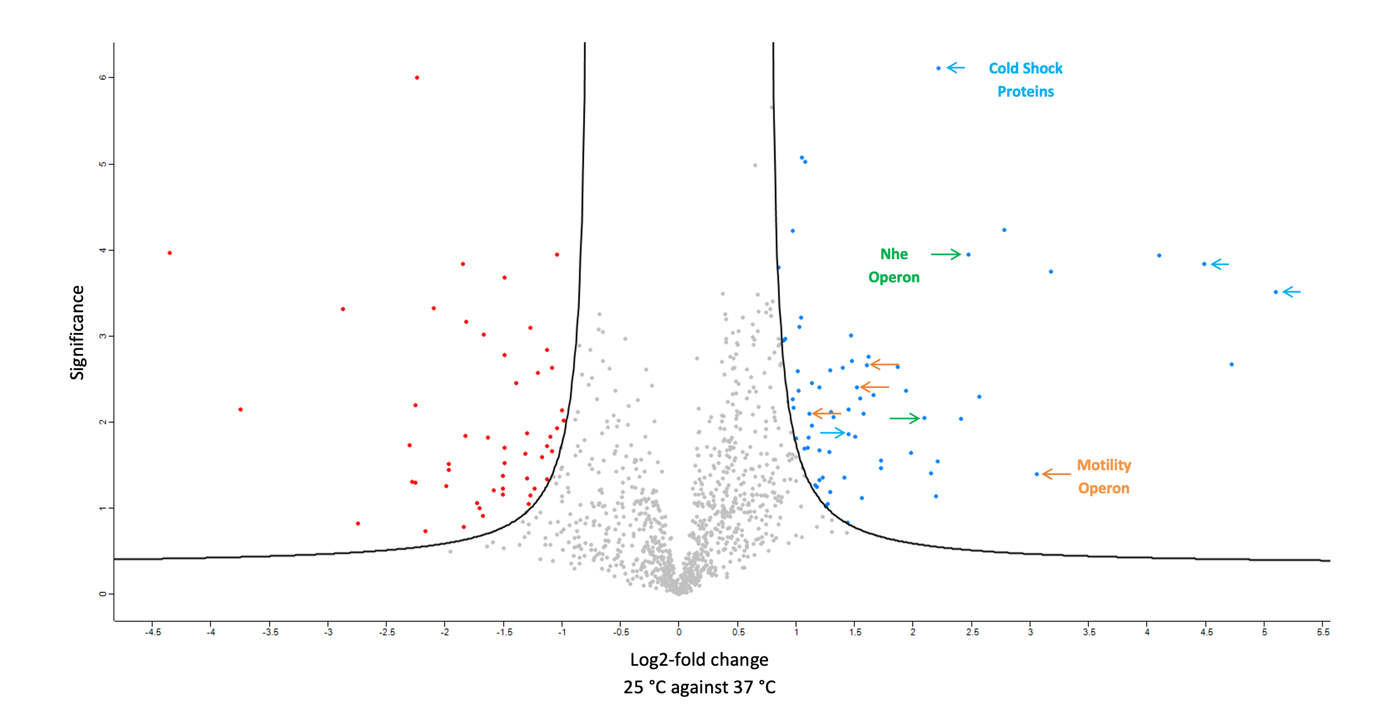

**Supplementary Figure S5. Temperature-dependent cell proteomes of mid-exponentially grown *Bc*G9241 cells**. Mass spectroscopy was used to analyse all the cellular proteins of *Bc*G9241, growing exponentially (OD_600_ = 0.5) at both 25°C and 37°C in LB broth, at 200 rpm. Peptide reads were counted using MaxQuant (Max Planck Institute) and comparisons were made with Perseus software (Max Planck Institute) and plotted as the difference in proteins expressed at 25°C against 37°C. Red and blue dots show proteins more highly secreted at 37°C and 25°C respectively. The black curved lines represent the significance cut off criteria of a P-value of <0.05 and a 2-fold change in protein level. Triplicate samples were analysed.

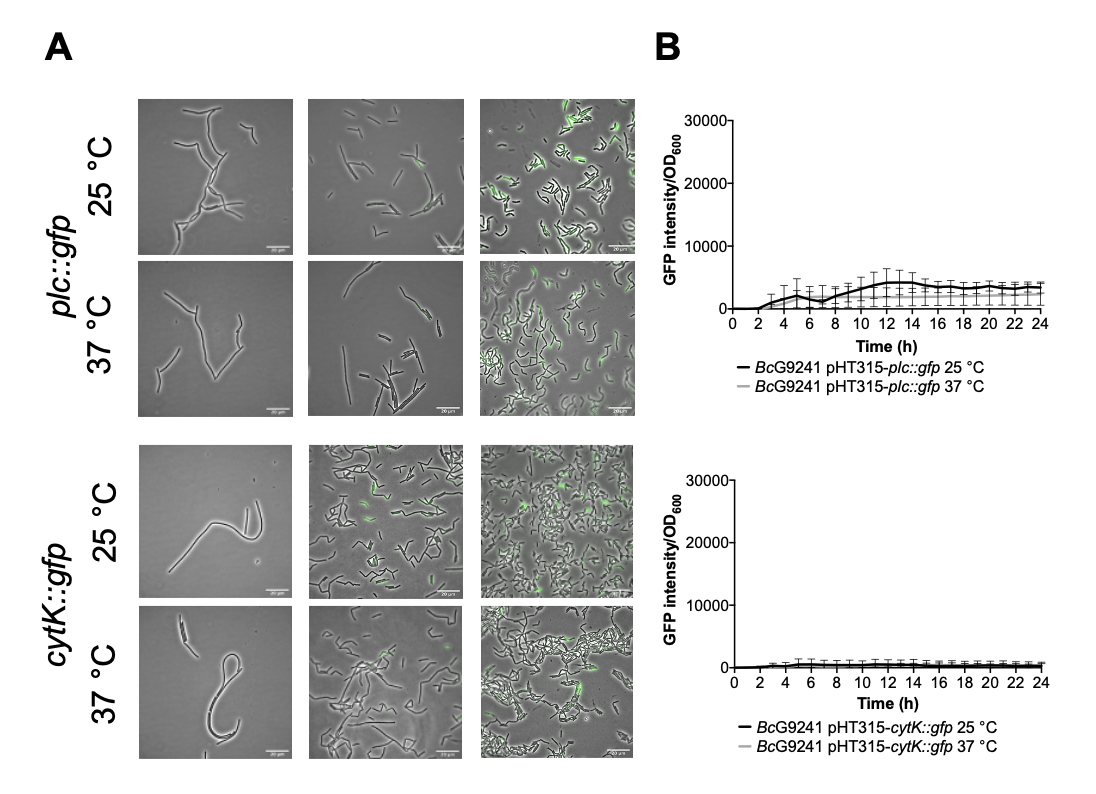

**Supplementary Figure S6**: **Temperature dependent expression of PlcR-regulated *plc* and *cytK* in *Bc*G9241 using GFP reporters.** (**A**) A representative selection of microscopy images of the transcription-translation GFP reporters of PlcR-regulated *plc* and *cytK* for *Bc*G9241 taken at three different time points: mid-exponential phase which is 2 hours at 37 °C and 5 hours at 25 °C (OD_600_=0.5), early stationary phase which is 4 hours at 37 °C and 7 hours at 25 °C (OD_600_=1.5) and 24 hours. (**B**) **Fluorescence of toxin reporters over time in LB.** GFP intensity/OD_600_ and change in GFP **(**$\boldsymbol{\Delta}$GFP/OD_600_) of *Bc*G9241 containing PlcR-regulated *plc* and *cytK* reporters over 24 hours growth in 100 µl volume LB media at 25 °C (in black) and 37 °C (in grey). Each line represents the mean of three biological replicates with three technical replicates each and error bars denote standard deviation.

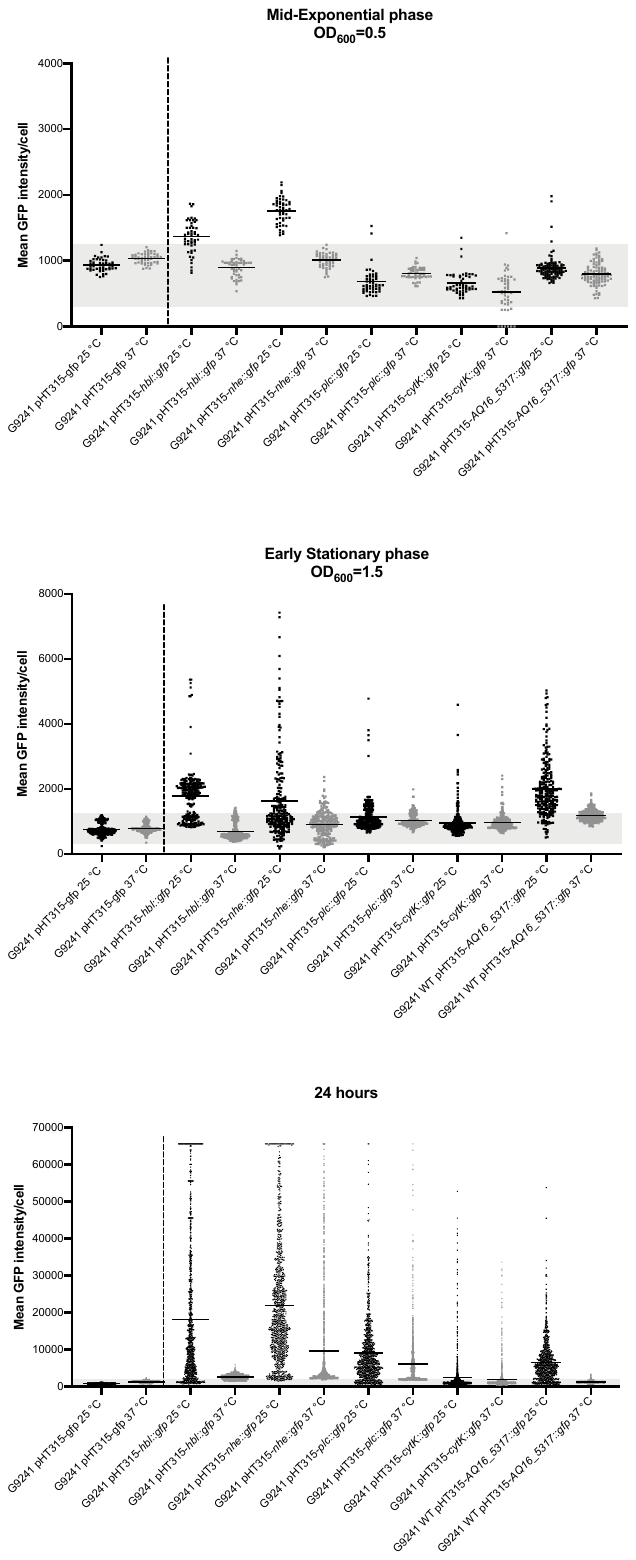

**Supplementary Figure S7: Quantification of GFP expression of PlcR-regulated proteins using image analysis.** Swarm-plots of GFP intensity of individual cells in the population of *Bc*G9241 at (**A**) mid-exponential (n = 50), (**B**) early stationary phase (n = 250) and (**C**) 24 hours (n = 1000) harbouring the transcription-translation toxin reporters compared to the pHT315-*gfp* no promoter control. The shaded area, selected by cells not expressing GFP, represents the threshold, where any dots above this area are cells expressing GFP. The line represents the mean of the overall GFP intensity.

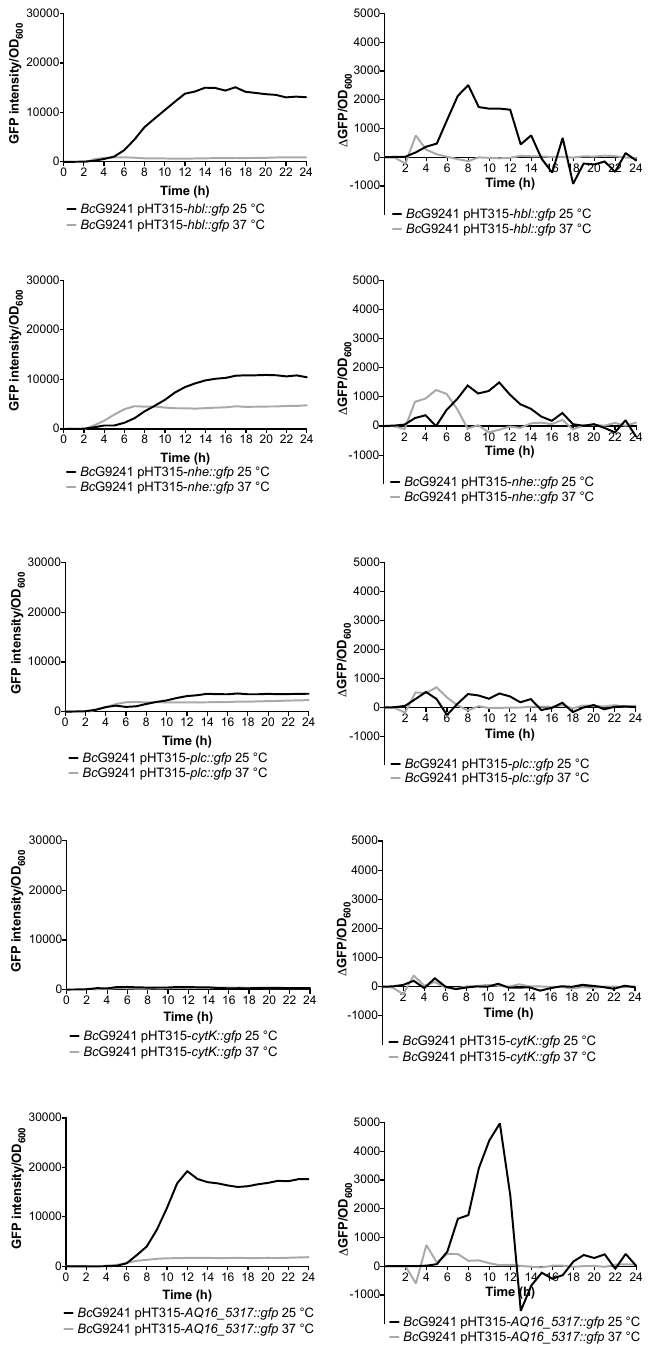

**Supplementary Figure S8: The rate of change in fluorescence from reporter strains in *Bc*G9241.** $\Delta$GFP/OD_600_ was calculated from this equation: $\Delta$GFP/OD_600_ = (GFP intensity_(t)_ - GFP intensity_(t-1)_)/OD_600._

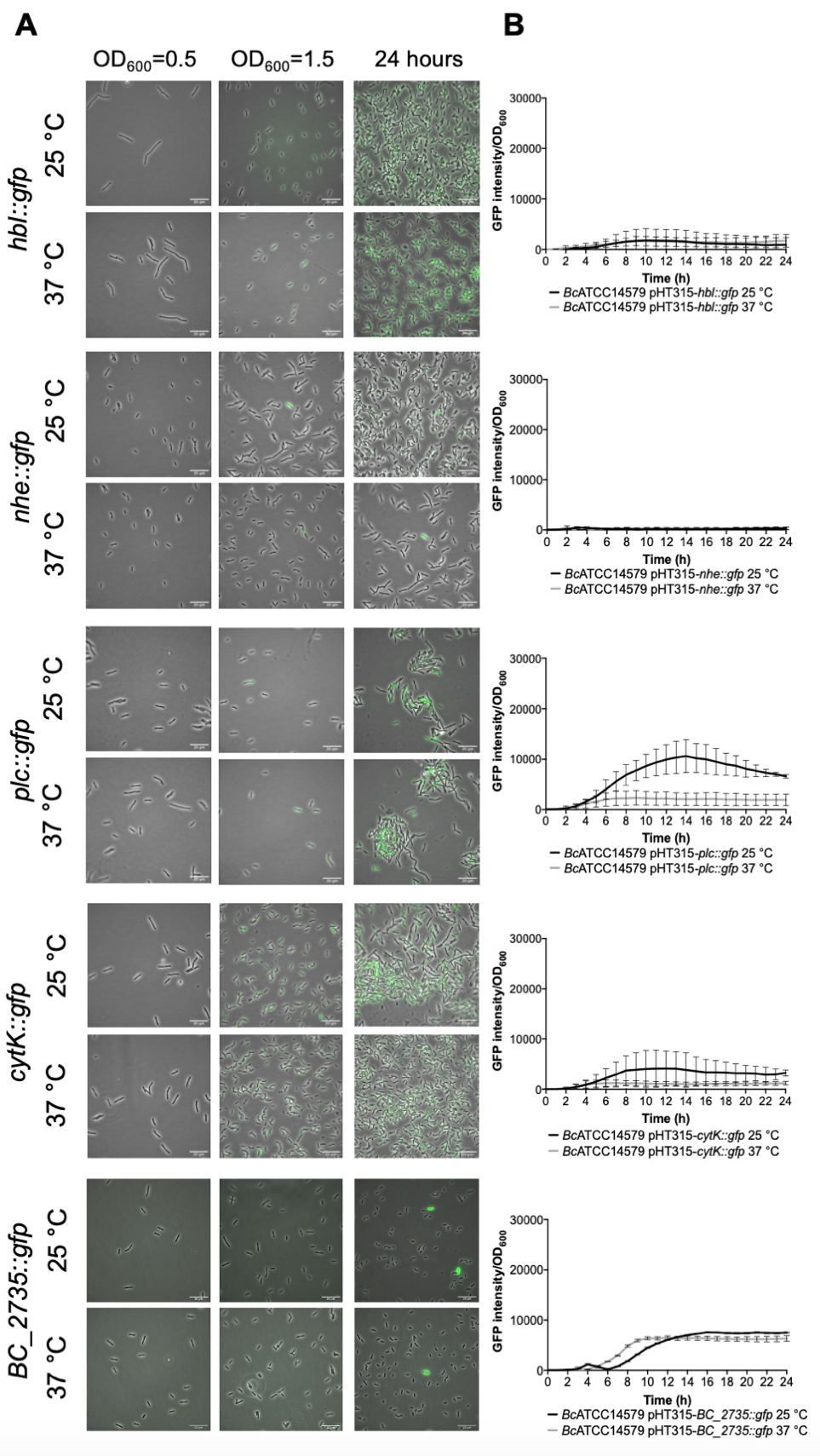

**Supplementary Figure S9: Expression of PlcR-regulated toxins and enzymes in *Bc*ATCC14579 using GFP reporters.** (**A**) A representative selection of microscopy images of the translational GFP reporters of PlcR-regulated toxins for *Bc*ATCC14579 taken at three different time points: mid-exponential phase which is 2 hours at 37 °C and 5 hours at 25 °C (OD_600_=0.5), early stationary phase which is 4 hours at 37 °C and 7 hours at 25 °C (OD_600_=1.5) and 24 hours. **(B)** GFP intensity/OD_600_ and change in GFP ($\Delta$GFP/OD_600_) of *Bc*ATCC14579 containing PlcR-regulated toxin reporters over 24 hours growth in 100 µl volume LB media at 25 °C (in black) and 37 °C (in grey). Each line represents the mean of three biological replicates with three technical replicates each and error bars denote standard deviation.

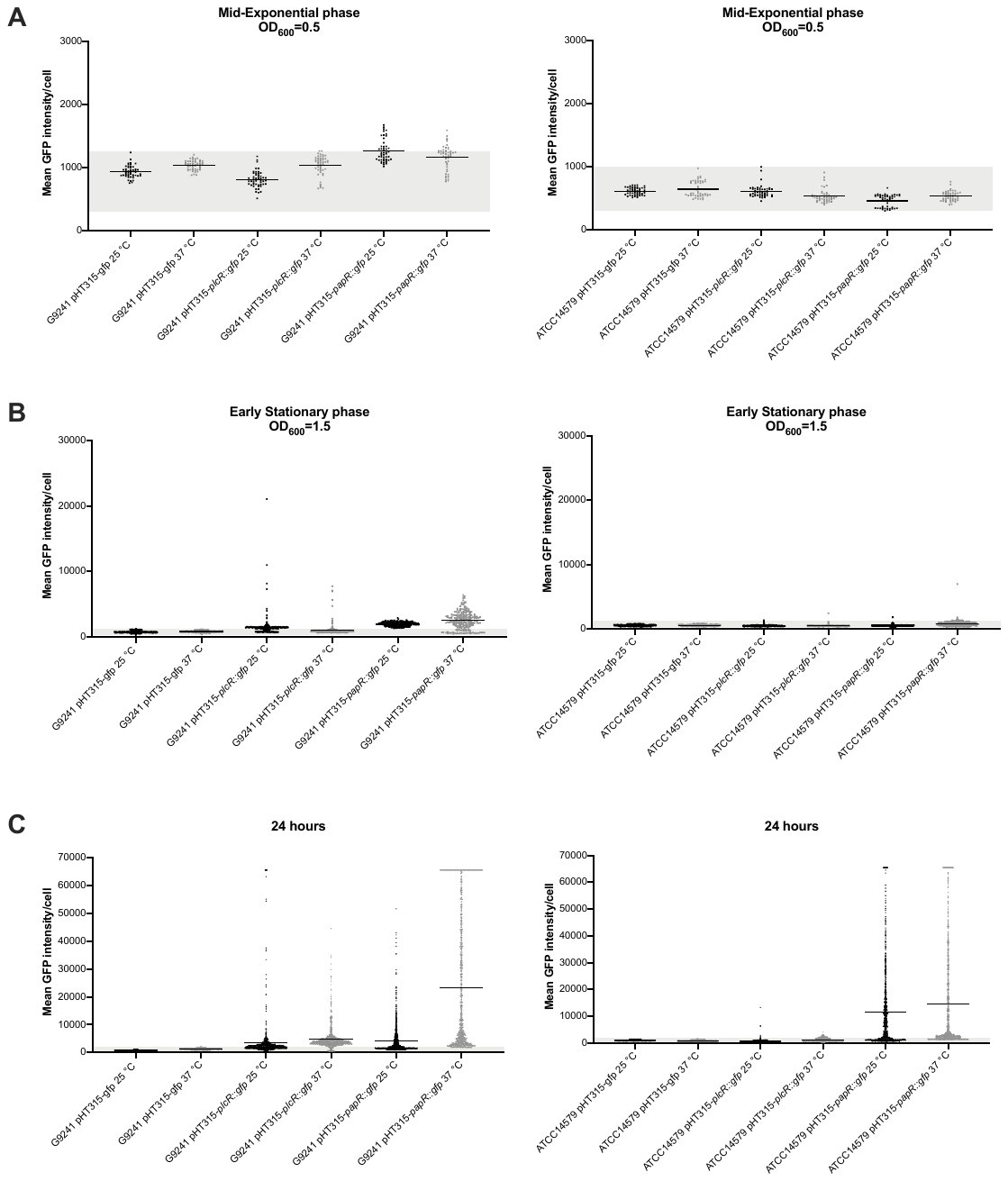

**Supplementary Figure S10: Quantification of PlcR and PapR expression using image analysis.** Swarm-plots of GFP intensity of individual cells in the population of *Bc*G9241 and *Bc*ATCC14579 harbouring pHT315-*plcR::gfp* or pHT315-*papR::gfp* compared to the pHT315-*gfp* no promoter control. GFP expression at (**A**) mid-exponential phase (N=50), (**B**) early stationary phase (N=250) and (**C**) 24 hours (N=1000). The line represents the mean of the overall GFP intensity. The shaded area, selected by cells not expressing GFP, represents the threshold, where any dots above this area are cells expressing GFP.

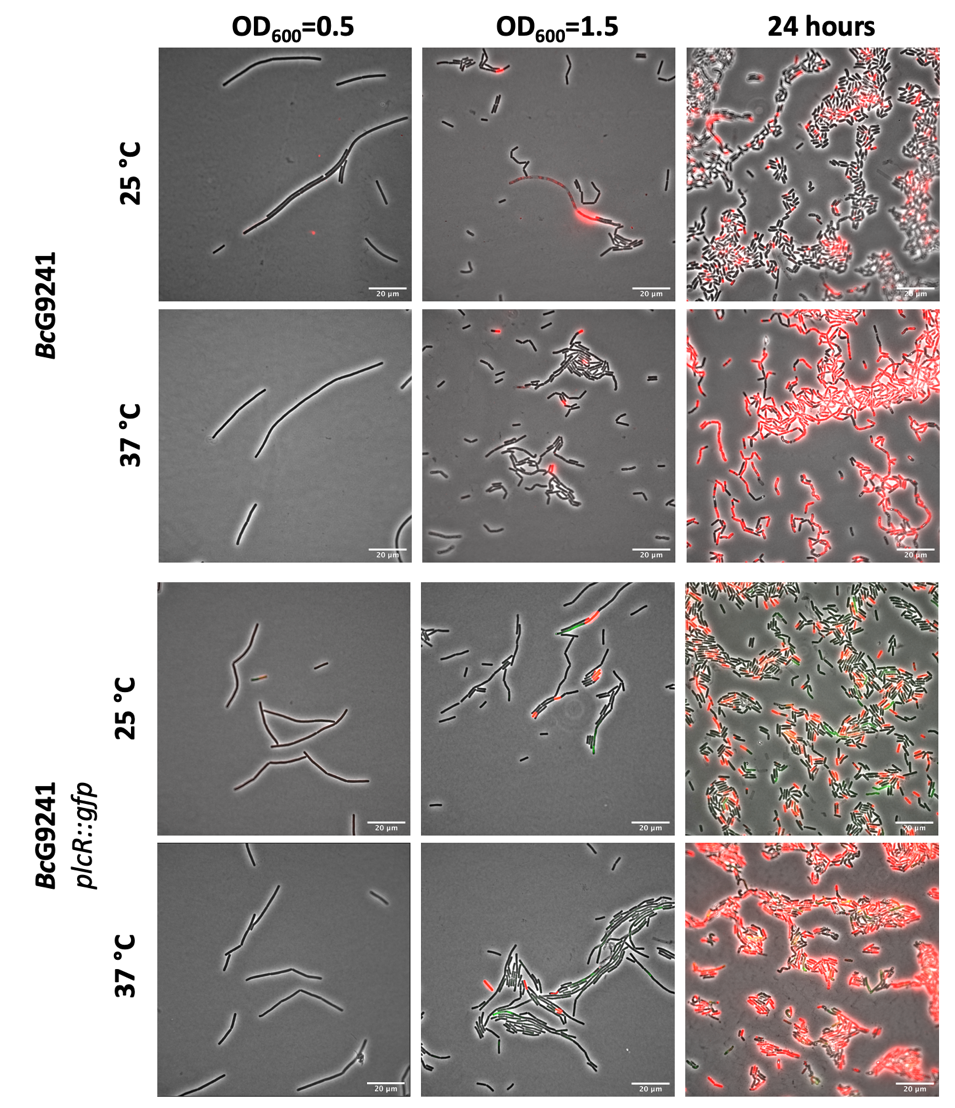

**Supplementary Figure S11: A representative selection of microscopy images of *Bc*G9241 stained with propidium iodide.** *Bc*G9241 WT and *Bc*G9241 harbouring the transcription-translation *plcR::gfp* reporter were stained with propidium iodide. Micrographs were taken at three different time points: mid-exponential phase which is 2 hours at 37 °C and 5 hours at 25 °C (OD_600_=0.5), early stationary phase which is 4 hours at 37 °C and 7 hours at 25 °C (OD_600_=1.5) and 24 hours. Scale bar = 20 𝜇m.

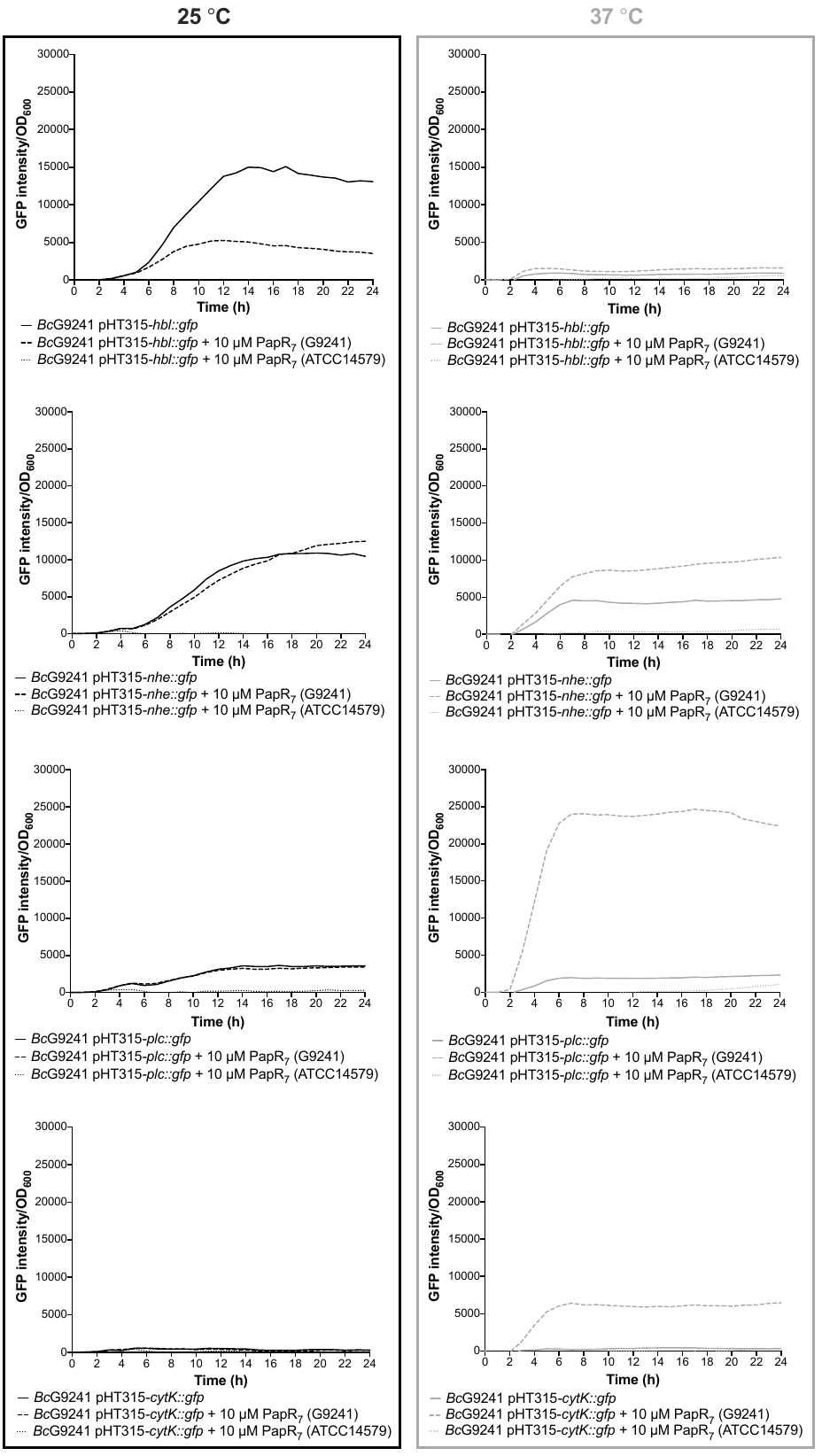

**Supplementary Figure S12: The effect of exogeneous PapR_7_ on the expression of PlcR-regulated toxins.** The overall GFP intensity/OD_600_ of the PlcR-regulated toxin reporters in *Bc*G9241, with and without the addition of synthetic PapR_7_, were measured over 24 hours in a plate reader at 25 and 37 °C. Dashed lines (--) represents the addition of G9241 PapR_7_ (SDLPFEH) and dotted lines (…) represents the addition of ATCC14579 PapR_7_ (KDLPFEY). All samples were to an n=3.

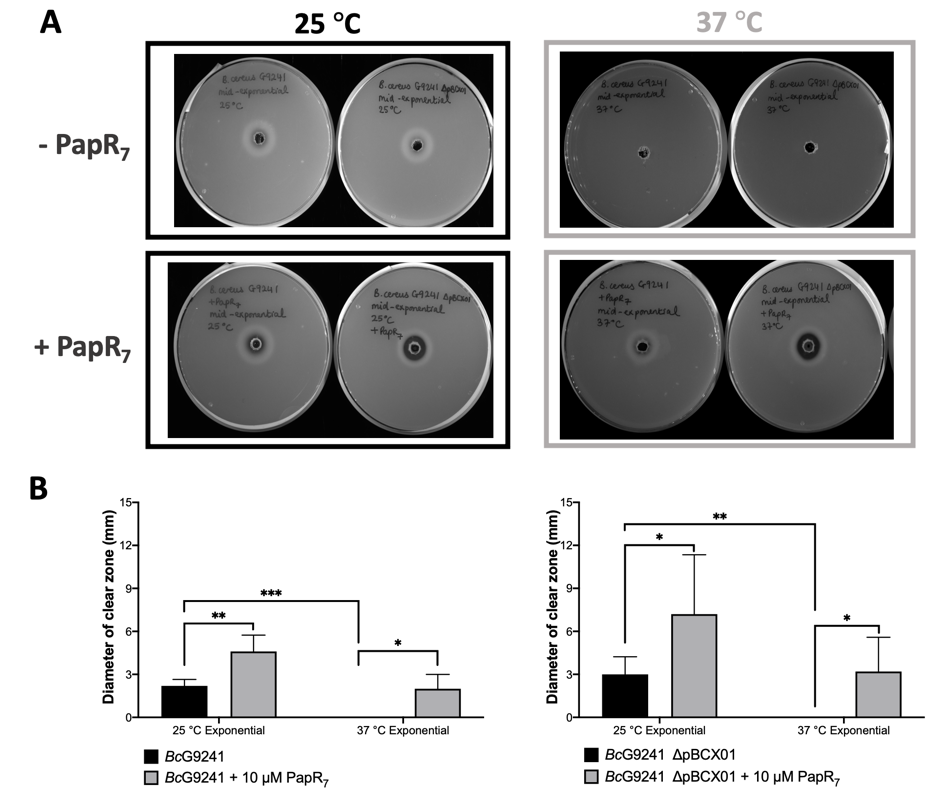

**Supplementary Figure S13: *Bc*G9241 expresses protease(s) in a temperature dependent manner**. (**A**) Hydrolysis zone of mid-exponential cell-free supernatants of *Bc*G9241 and *Bc*G9241 $\Delta$pBCX01 on skim milk agar plates observed at 25 $^{\circ}$C but not at 37 $^{\circ}$C. (**B**) Diameter (mm) of clear zone produced by secreted proteases on LB + 2% (w/v) skim milk agar plates for 24 hours of incubation at 37 $^{\circ}$C. Values are represented as mean and standard deviation, number of replicates, n = 5. * [P < 0.05], **[P < 0.01] and ***[P < 0.001] as determined by paired t-test.

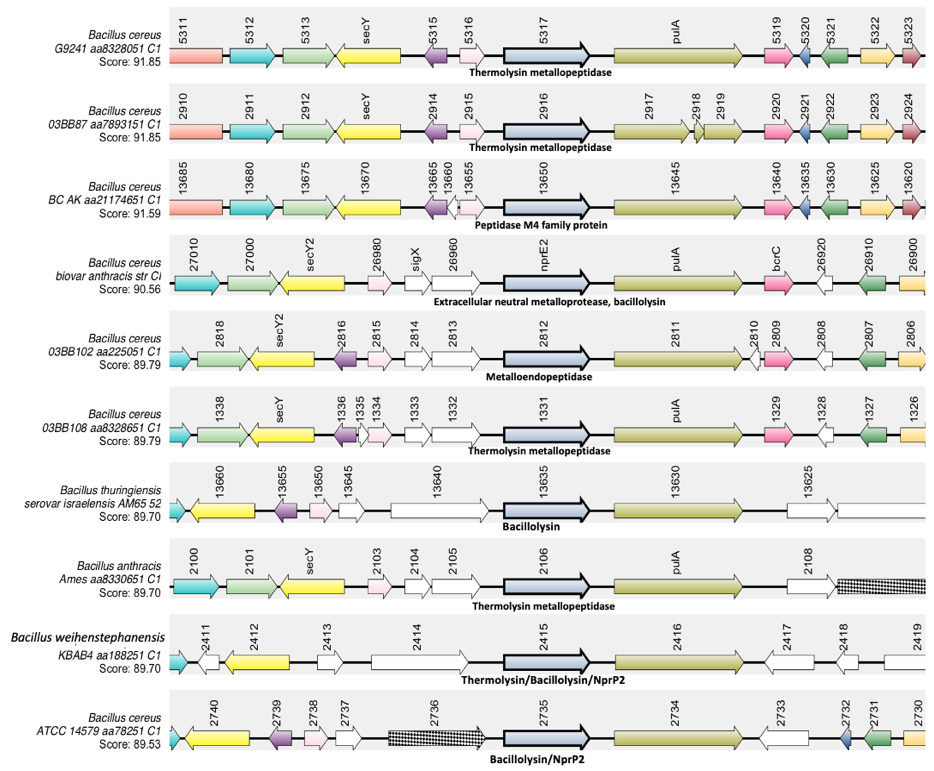

**Supplementary Figure S14: Synteny of the gene encoding thermolysin metallopeptidase in *B. cereus* sensu stricto, *B. thuringiensis, B. weihenstephanensis, B. anthracis* and *B. cereus-B. anthracis* crossover strains*.*** The protein sequence of AQ16_5317 from *Bc*G9241 (RefSeq accession GCF_000832815.1) was used as the query protein. The gene AQ16_5317 and its homologs are shown in blue in bold. SynTax, a synteny web service, was used to look in the conservation of gene order (<https://archaea.i2bc.paris-saclay.fr/SyntTax/Default.aspx>).

**Supplementary Tables**

**Supplementary Table S1: Toxin genes found in *Bc*G9241** **with** **PlcR-box motif identified upstream of the gene (highlighted grey).** Subscript numbers denote the distance in bp upstream of gene.

| **Gene** | **Upstream of Loci** | **Sequence** |
| --- | --- | --- |
| *nhe* | AQ16_660 – 658 | _550_TTTGTATACACTATGCATAATTGCATATGAGTCCAAAA_509_ |
| *hbl* | AQ16_4930 – 4933 | _907_TATCTACATTTTATGCAATTATACATAACTAAATAAAG_866_ |
| Collagenase | AQ16_1941 | _95_AGAAGAAATAATATGAAATATTGCATTTTATATTGTTG_56_ |
| Thermolysin | AQ16_5317 | _123_CGTCCTTATATTATGTAATTTTGCATAATGTTACATAA_86_ |
| *plc* | AQ16_1823 | _150_AGTTATAATGATATGAACATTTGCATATTTTAATTTAG_113_ |
| *cytK* | AQ16_1392 | _112_CAAAACTCACCTATGCAATTATGCATAACTATCCCTTC_75_ |

**Supplementary Table S2: Plasmids used in this study.**

| **Plasmid name** | **Relevant properties** | **Reference** |
| --- | --- | --- |
| pHT315 | Bears the replication region of the *B. thuringiensis* pHT1030. ori from pBR322, erythromycin resistance gene, ampicillin resistance gene, pUC19 base. 15 ± 5 copies/chromosome. | (62) |
| pHT315-*gfp* | gfp ORF cloned into pHT315. | This study |
| pHT315-*plcR::gfp* | PlcR transcription-translation reporter. pHT315-gfp with *plcR* promoter and eight amino acids from the PlcR ORF fused with *gfp*. | This study. |
| pHT315-*papR::gfp* | PapR transcription-translation reporter. pHT315-gfp with papR promoter and eight amino acids from the PapR ORF fused with gfp. | This study. |
| pHT315-*hbl::gfp* | Hbl transcription-translation reporter. pHT315-gfp with hbl promoter and eight amino acids from the Hbl ORF fused with gfp. | This study. |
| pHT315-*nhe::gfp* | Nhe transcription-translation reporter. pHT315-gfp with nhe promoter and eight amino acids from the Nhe ORF fused with gfp. | This study. |
| pHT315-*plc::gfp* | Plc transcription-translation reporter. pHT315-gfp with plc promoter and eight amino acids from the Plc ORF fused with gfp. | This study. |
| pHT315-*cytK::gfp* | CytK transcription-translation reporter. pHT315-gfp with cytK promoter and eight amino acids from the CytK ORF fused with *gfp.* | This study. |
| pHT315-*AQ16_5317::gfp* | Thermolysin metallopeptidase transcription-translation reporter. pHT315-*gfp* with AQ16_5317 promoter (including the native Shine Dalgarno region) and eight amino acid of the open reading frame directly upstream of *gfp* | This study. |
| pHT315-*nprB* | pHT315 with the promoter and the full open reading frame of *nprB* (amplified from *B. cereus* ATCC14579) | This study. |
| pHT315-*fusA_p_* | pHT315 with *fusA* promoter (including the native Shine Dalgarno region) and the full open reading frame of AQ16_5317 | This study. |
| pHT315-*fusA_p_-AQ16_5317* | pHT315 with *fusA* promoter (including the native Shine Dalgarno region) and the full open reading frame of AQ16_5317 | This study. |

**Supplementary Table S3: Primers used in this study.** F=Forward, R=Reverse, G9241=Primers used for *B. cereus* G9241, ATCC=Primers used for *B. cereus* ATCC14579. Primers were synthesised and supplied by Integrated DNA Technologies (IDT).

| **PRIMER No** | **PRIMER DESCRIPTION** | | **PRIMER 5’ – 3’** | | **Restriction**  **enzyme** |
| --- | --- | --- | --- | --- | --- |
| To insert *gfp* in pHT315 | | | | | |
| 1 | GFP_F | | GATATAggtaccGCTAGCAAAGGAGAA | | *Kpn*I |
| 2 | GFP_R | | CCGAGCTCgaattcATTATTTGTAGA | | *Eco*RI |
| **TRANSCRIPTION-TRANSLATION FUSION PlcR and PapR**  Vector used to insert amplified fragment: pHT315-*gfp* | | | | | |
| 3 | G9241_plcR_F | | ATAcTgcAGACCGAATGTAACGAAAGCATAGGCAA | | *Pst*I |
| 4 | G9241_plcR_translate_R | | CTTggtaccACTTCCTAATTTTTCTGCGTGCAT | | *Kpn*I |
| 5 | G9241_papR_F | | TATcTgcAGTGGTATATTTTGTAGGTTATATCCA | | *Pst*I |
| 6 | G9241_papR_translate_R | | CGTggtaccACTACCAATAAGTAATTTTTTCAT | | *Kpn*I |
| 7 | ATCC_plcR_F | | ACACTgCagATCTATTATTGTATGTGAGATGAA | | *Pst*I |
| 8 | ATCC_plcR_translate_R | | CTTggtaccACTTCCTAATTTTTCTGCGTGCAT | | *Kpn*I |
| 9 | ATCC_papR_F | | TAActgCAGTAGATAAATATCCTAAAAAATGGGTA | | *Pst*I |
| 10 | ATCC_papR_translate_R | | CGTggtaccACTACCAATAAGTAATTTCTTCAT | | *Kpn*I |
| **TRANSCRIPTION-TRANSLATION FUSION PlcR-regulated toxins**  Vector used to insert amplified fragment: pHT315-*gfp* | | | | | |
| 11 | F_G9241_cytK_translate | | ATAcTgcAGATTCCCTTGAGAAGGATT | | *Pst*I |
| 12 | R_ G9241_cytK_translate | | CGTggtaccACGTTTTAGAGCGTTTCAT | | *Kpn*I |
| 13 | F_ G9241_plc_translate | | ATAcTgcAGTTTAGACAAGCCTTAATA | | *Pst*I |
| 14 | R_ G9241_plc_translate | | CTTggtaccAAGTACTTTCTTTTTCAT | | *Kpn*I |
| 15 | F_ G9241_nhe_translate | | ATAcTgcAGAAGGATAGCTTTACGAAA | | *Pst*I |
| 16 | R_ G9241_nhe_translate | | CTTggtaccAATTAAAGTCTTTTTCAC | | *Kpn*I |
| 17 | F_ G9241_hbl_translate | | CCAcTgcAGATGTAAAGACTTAAACAG | | *Pst*I |
| 18 | R_ G9241_hbl_translate | | CTTggtaccCATTATTTTAGTTTTCAT | | *Kpn*I |
| 19 | F_ATCC_cytK_translate | | ATActgcagCAATCTTCAGAATAAGAA | | *Pst*I |
| 20 | R_ATCC_cytK_translate | | CGTggtaccGGTTTTAGAACGTTTCAT | | *Kpn*I |
| 21 | F_ATCC_plc_translate | | ATActgcagCATTAGTTTAGACAAGCC | | *Pst*I |
| 22 | R_ATCC_plc_translate | | CGTggtaccAGCAAGTACTTTTTTTTTCAT | | *Kpn*I |
| 23 | F_ATCC_nhe_translate | | CTActgcagCATATTTACGCATGTTGT | | *Pst*I |
| 24 | R_ATCC_nhe_translate | | CGCggtaccAATTAAAGTCTTTTTCAC | | *Kpn*I |
| 25 | F_ATCC_hbl_translate | | ATActgcagCATGGGTATAAGTATCCC | | *Pst*I |
| 26 | R_ATCC_hbl_translate | | CGTggtaccTCCTGTAATTATTTTAGTTTTCAT | | *Kpn*I |
| 27 | F_G9241_AQ16_5317_translate | | ATActgcagATGTTTCTTTCTAGCATT | | *Pst*I |
| 28 | R_G9241_AQ16_5317_translate | | ATAggtaccCGTCTTTTTGTTTTTCAT | | *Kpn*I |
| 29 | F_ATCC_BC_2735_translate | | GCGctgcagTTATAAAATTTCTTATAG | | *Pst*I |
| 30 | R_ATCC_BC_2735_translate | | ATAggtaccTGTCTTTTTGTTTTTCAT | | *Kpn*I |
| **AQ16_5317 under *fusA* promoter**  Vectors used to insert amplified fragment: pHT315 and pHT315-*fusAp* | | | | | |
| 31 | F_G9241_fusA_promoter | | AATctgcagGTGAAGACACTCATAAAATGGC | | *Pst*I |
| 32 | R_G9241_fusA_promoter | | ATTggatccCATTTGGTGTCTTGCTCCTT | | *Bam*HI |
| 33 | F_G9241_AQ16_5317 | | CGAggatccAAAAACAAAAAGACGTTAAC | | *Bam*HI |
| 34 | R_G9241_AQ16_5317 | | GCCggtaccTTACTTAATTTTTGCTGCAT | | *Kpn*I |

**Supplementary Materials and Methods**

**Cell cultures.** T2 cells, an immortalized T-cell line (63), were grown in RPMI-1640 (Sigma) supplemented with 10% foetal bovine serum (Gibco), 1 mM L-glutamine, 200 units/ml penicillin, and 0.1 mg/mL streptomycin (Sigma) at 37 °C in a 5% CO_2_ atmosphere. Polymorphonuclear leukocytes (PMNs) were isolated from freshly collected human blood obtained from volunteers as previously described (60). Differentiation of macrophages from THP-1 human monocytic cell line was done by induction with phorbol myristate acetate (PMA), following the protocol described by (64).

**Cytolytic activity assays**. To evaluate the cytotoxicity of culture supernatants, strains were grown at 25 ºC and 37 ºC in LB broth. The cultures were recovered after 16 h and normalized to an optical density at 600 nm (OD_600_) of 1. Cells were pelleted by centrifugation (4000 *g* for 10 min) and the supernatants were sterilized by filtration through 0.22 µm pore size filters (Millipore).

**(i)** To evaluate the lytic activity of *Bacillus* supernatants to haemocytes, haemolymph from *M. sexta* larvae were collected as previously described (65). Haemocytes in Grace’s medium (100 µL) were added to a 96-well flat-bottom polystyrene plate and incubated with bacterial supernatant (50 µL) for 1 h at room temperature. Reaction mixture was removed and attached cells were rinsed and resuspended in Grace’s medium to obtain a suspension of 1×10^5^ to 1×10^6^ cells/mL. Cell viability was measured using the Guava EasyCyte™ flow cytometry and Guava® ViaCount® Reagent (Millipore), following the manufacturer’s instructions. Assays were done by triplicate. For observation under microscopy, the final suspension cell was incubated with acridine orange (10 μg/mL) for 2 min and then viewed and imaged with filters for green fluorescence in a laser-scanning confocal microscope (Zeiss).

**(ii)** Cytotoxicity towards immune mammalian cells were tested using T2 cells, PMNs, and macrophages differentiated from THP-1 human monocytes. Cell suspensions in RPMI-1640 medium (100 µl) were incubated with 50 μL of bacterial supernatant in a 96-well flat-bottom polystyrene plate for 90 minutes at 37 ºC. Reaction mixture was removed and attached cells were rinsed and resuspended in PBS to obtain a suspension of 1×10^5^ to 1×10^6^ cells/mL. Cell viability was assayed by using Guava® ViaCount® Reagent and Guava EasyCyte™ flow cytometry, following the manufacturer’s instructions. Assays for each cell type were done by triplicate.

**Creation of electro-competent *E. coli* ET12567 and transformation.** 500 ml of *E. coli* were grown to OD_600_=0.5 and cells were harvested by centrifugation at 5000 rpm for 15 minutes at 4 °C. Cells were washed first with 300 ml of 4 °C sterile H_2_O, then with 50 ml 10% (v/v) glycerol at 4 °C and finally resuspended in 1ml 10% (v/v) glycerol. 50 µl aliquots were stored at -80 °C. 80 µl of electrocompetent *E. coli* ET12567 cells were combined with 2 µl of plasmid in a pre-chilled Gene Pulser®/Micropulser™ electroporation cuvette, 2 mm (Bio-Rad). Cells were pulsed 2.5 kV, 25 µF and 200 Ω. 1 ml ice cold LB broth was added to cells after electroporation and bacteria were incubated at 37 °C for 1 hour to recover. Cells were plated onto selective media and successful transformants isolated.

**Transformation of *B. cereus* species.** Overnight cultures were diluted 1:100 in 50 ml of LB medium. Cells were cultured at 37 °C and harvested at OD_600_ of 0.5 by centrifugation at 5000 g for 10 minutes at 4˚C. Cell pellets were resuspended in 10 ml of 10% glycerol in ddH_2_O. The wash was repeated two more times. Washed cell pellets were resuspended in 100 µl of ddH_2_O to create ~220 µl of competent cells. 2 µg of plasmid DNA was added to the cells and 50 µl of this cell-plasmid mix was aliquoted into a pre-chilled 2 mm cuvette. Cells were electroporated at 2.5 kV, 25 µF and 200 Ω. 1 ml of SOC media (NEB) was immediately added to aid cell recovery. Cells were transferred to a 15 ml falcon tube and incubated at 37 °C for 3 hours. Transformants were selected on the appropriate media.

**Image analysis of reporter strain expression.** Quantitative image analysis of micrographs of *Bc*G9241 and *Bc*ATCC14579 harbouring GFP reporters was performed using Fiji, as described by (66). Brightfield images were used to automatically detect the bacteria by converting the image to binary, followed by edge detection and particle analysis with the size of the particles set to 0.5–10 μm^2^ for rod shapes and 0.5–200 μm^2^ when the bacteria are filamenting. The resulting regions of interest were then used to measure the intensity in the corresponding green channel (for GFP detection). At least four images were used per time point with a minimum total number of cells of 50 for each time point. The control consists of images taken of *Bc*G9241 and *Bc*ATCC14579 harbouring the no promoter control plasmid pHT315-*gfp* and for the purposes of this analysis the mean intensity per cell calculated for each time point were combined to find a threshold value for autofluorescence.

**Protease activity assay.** Secreted proteins were collected from mid-exponential (OD_600_ ~0.5) cultures grown at both 25 °C and 37 °C, with or without the addition of PapR_7_ during lag growth phase. 15 ml of supernatant was extracted (5 minutes at 6,000 x g) and filtered (0.22 $\mu$m) before being concentrated using the 3K Amicon® Ultra Centrifugal Filters (Millipore) to 50 μL. Screening for protease expression of *Bc*G9241 grown at 25 °C and 37 °C was carried out using LB agar containing 2% (w/v) skim milk plates. Plates were incubated at 37 °C for 24 hours, and clearing halos were determined.
